# The COMPASS-like complex modulates fungal development and pathogenesis by regulating H3K4me3-mediated targeted gene expression in *Magnaporthe oryzae*

**DOI:** 10.1101/2020.03.13.990218

**Authors:** Sida Zhou, Wanyu Sun, Xinyu Zhao, Yang Xu, Mengyu Zhang, Yue Yin, Song Pan, Dan He, Mi Shen, Jun Yang, Xiuying Liu, Qi Zheng, Weixiang Wang

## Abstract

Histone H3K4 methylation is catalysed by the multi-protein complex known as the Set1/COMPASS or MLL/COMPASS-like complex, an element that is highly evolutionarily conserved from yeast to humans. However, the components and mechanisms by which the COMPASS-like complex targets the H3K4 methylation of plant pathogenic genes in fungi remain elusive. Here we present a comprehensive analysis combining biochemical, molecular, and genome-wide approaches to characterize the roles of the COMPASS-like family in *Magnaporthe oryzae*, a model plant fungal pathogen. We purified and identified six conserved subunits of COMPASS from the rice blast fungus *M. oryzae*, i.e., MoBre2 (Cps60/ASH2L), MoSpp1 (Cps40/Cfp1), MoSwd2 (Cps35), MoSdc1 (Cps25/DPY30), MoSet1 (MLL/ALL) and MoRbBP5 (Cps50), using an affinity tag on MoBre2. We determined the SPRY domain of MoBre2 can recognize directly with DPY30 domain of MoSdc1 *in vitro.* Furthermore, we found that deletion of the genes encoding COMPASS subunits of MoBre2, MoSpp1 and MoSwd2 caused similar defects regarding invasive hyphal development and pathogenicity. Genome-wide profiling of H3K4me3 revealed that the it has remarkable co-occupancy at the TSS regions of target genes. Significantly, these target genes are often involved in spore germination and pathogenesis. Decreased gene expression caused by the deletion of *MoBre2*, *MoSwd2* or *MoSpp1* gene was highly correlated with decrease in H3K4me3. Taken together, these results suggest that MoBre2, MoSpp1, and MoSwd2 function as a whole COMPASS complex, contributing to fungal development and pathogenesis by regulating H3K4me3-targeted genes in *M. oryzae*.

## INTRODUCTION

The eukaryotic genome is found in complexes with an equal mass of histone proteins to form nucleosomes, the basic units of chromatin (Kornberg, 1974; Smith, 2010). Chromatin assembly and folding into higher structures is a dynamic process that ultimately affects the activation or repression of gene expression. Chromatin remodelling and covalent modifications of histones are two means by which variation is introduced into the chromatin polymer, thereby regulating the structure and function of chromatin and ultimately gene expression (Luger et al., 1997; Strath, 2000; Lachner, 2002; Bhaumik et al., 2007; Shilatifard, 2008). Chromatin-modifying activities are often recruited to specific gene regulatory sequences, whereupon they cause localized changes in the chromatin structure and specific transcriptional effects (Ng et al., 2003; Allis, 2007). Several post-translational modifications are known to be present on the terminal domains of histones, such as acetylation, phosphorylation, ubiquitination, ADP ribosylation, and methylation (Workman et al., 2003; Berger et al., 2007; Kouzarides, 2007). Lysine methylation has been well documented on histones H3 and H4 *in vivo*, and residues 4, 9, 27, and 36 are typical methylation sites (Wang et al., 2010; Yang et al., 2012).

Histone H3K4 methylation is associated with active chromatin in a wide range of eukaryotic organisms. Many of the genes involved in H3K4 methylation encode proteins bearing a 130- to 140-amino-acid motif called the SET domain. This domain takes its name from the *Drosophila* proteins *<u>S</u>u(var)3-9*, *<u>E</u>nhancer of zeste (E(z))* and *<u>T</u>rithorax*. Many SET domain-containing proteins have been shown to possess histone or lysine methyltransferase activity (Tschiersch et al., 1994; Stassen et al., 1995; Shilatifard, 2012). In *Saccharomyces cerevisiae*, the Set1/COMPASS (<u>C</u>omplex <u>P</u>roteins <u>A</u>ssociated with <u>S</u>et1), which consists of the seven polypeptides Set1, Cps60/Bre2, Cps50/Swd1, Cps40/Spp1, Cps35/Swd2, Cps30/Swd3, and Cps25/Sdc1, was the first H3K4 methylase to be identified (Miller et al., 2001; Krogan et al., 2003). Cps60 is the product of the *BRE2/YLR015w* gene, which encodes a protein similar to both *Drosophila* and human ASH2L proteins. ASH2L is a member of the Trx family of homeodomain DNA-binding proteins that is thought to regulate gene expression, morphogenesis, and differentiation in humans. Cps25 is encoded by the *YDR469* gene, which has weak similarity to a DPY30 protein involved in dosage compensation of expression of genes on the X chromosome. Some studies have revealed that Cps60 (Bre2) and Cps25 (Sdc1) form a globular structure sitting at the base of the Y-shaped COMPASS structure sandwich (Chen et al., 2012; Li et al., 2016). Cps35 is the only essential subunit of the COMPASS. In human embryonic kidney 293 cells, the depletion of WDR5, (a Cps35 homologue) can result in a global loss of histone H3K4 methylation and concomitant loss of Hox gene expression (Li et al., 2016). The *mammalian* Wdr82 (a Cps35 homologue) is required for the targeting of Setd1A-mediated histone H3-Lys4 trimethylation near transcription start sites via tethering to RNA polymerase II, an event that is a consequence of transcription initiation (Lee et al., 2008; Palmer et al., 2013). Null mutants missing any one of the six COMPASS subunits grow more slowly and display greater hydroxyurea sensitivity than wild-type cells under normal conditions and they also exhibit a defect in silencing gene expression near chromosomal telomeres (Roguev et al., 2001; Dover et al., 2002).

Previous studies have shown that the COMPASS family of H3K4 methyltransferases is highly conserved from yeast to plants and humans (Jiang et al., 2011; Shilatifard, 2012). In *Drosophila melanogaster*, there are three subclasses within COMPASS-like complexes, namely SetD1A, Trx and trithorax-related (Trr), all of which are homologues to yeast Set1 (Ardehali et al., 2011; Mohan et al., 2011; Shilatifard, 2012). While Set1D1A is homologous to yeast Set1, Trx and Trr are more distantly related (Czermin et al., 2002). dSet1 (*Drosophila Set1*), Trx and Trr each interact with a set of core complex subunits, with additional subunits unique to each (Mohan et al., 2011). In human cells, Set1A, Set1B, MLL1, MLL2, MLL3 and MLL4 are homologous to yeast Set1 (Cho et al, 2007). A biochemical study showed that human cells bear at least six COMPASS family members, each of which is capable of methylating H3K4 with non-redundant functions (Roguev et al., 2001; Dover 2002). These enzymes participate in diverse gene regulatory networks and are fully active in the context of an MLL/COMPASS-like complex that includes the proteins: WDR5, RbBP5, ASH2L and DPY-30 (WRAD). WRAD components function by engaging in physical interactions that recruit the SET1 family of proteins to target chromatin sites (Ernst et al., 2012; Ali et al., 2017; Hsu et al., 2018).

Recently, in filamentous fungi, such as *Fusarium spp*. and *Aspergillus spp*., H3K4 methylation is also associated with active transcription of the majority of genes. Deletion of the COMPASS components CCLA and CCL1 in *Aspergillus spp*. and *Fusarium spp*., respectively, results in increased expression of several, secondary metabolites (SMs) in these fungi (Palmer et al., 2013; Shinohara et al., 2016; Studt et al., 2016). Some studies have shown that H3K4 methylation is also associated with mainly secondary metabolism cluster genes, in the filamentous fungi, such as *Asperigillus spp.* and *Fusarium spp*. (Connolly et al., 2013; Palmer et al., 2013; Matthews et al., 2016). In *F. graminearum*, *FgSet1* is a Set1 orthologue and is one subunit of the COMPASS-like complex, which also plays an important role in regulating fungal growth and secondary metabolism (Connolly et al., 2013; Liu et al., 2015; Bachleitner et al., 2019).

*M. oryzae* functions as a model plant pathogenic fungus as the causal agent of rice blast disease, which is the most devastating and persistent disease of cultivated rice (Cao et al., 2016; Yan, 2016; Liu et al., 2018; Zhang et al., 2018). MoSet1 is the homologue of *S. cerevisiae* Set1. When the *MoSet1* is depleted (Δ*moset1*), H3K4 methylation becomes impaired, resulting in severe defects in infection-related morphogenesis, including in conidiation and appressorium formation (Pham et al., 2015). The homologues of COMPASS complex exit in multiple fungi, but other than the Set1 subunit, no clear systemic functions of COMPASS have been reported in *M. oryzae*.

Here, we report the identification of a conserved COMPASS-like complex that contains six subunits: MoSet1, MoBre2, MoSwd1, MoSpp1, MoSwd2 and MoSdc1(Table 1). When the component genes, *MoBre2*, *MoSpp1* and *MoSwd2* were deleted, H3K4me3 signals were dramatically decreased. Our results show that MoBre2, MoSpp1 and MoSwd2 not only have unique regulation roles in fungal development, but also function as a whole complex to methylate H3K4 in the TSS region of its target genes. Moreover, most of these key target genes are involved in conidia, mycelial growth and infectious hyphal formation in *M. oryzae*.

**Table 1.**
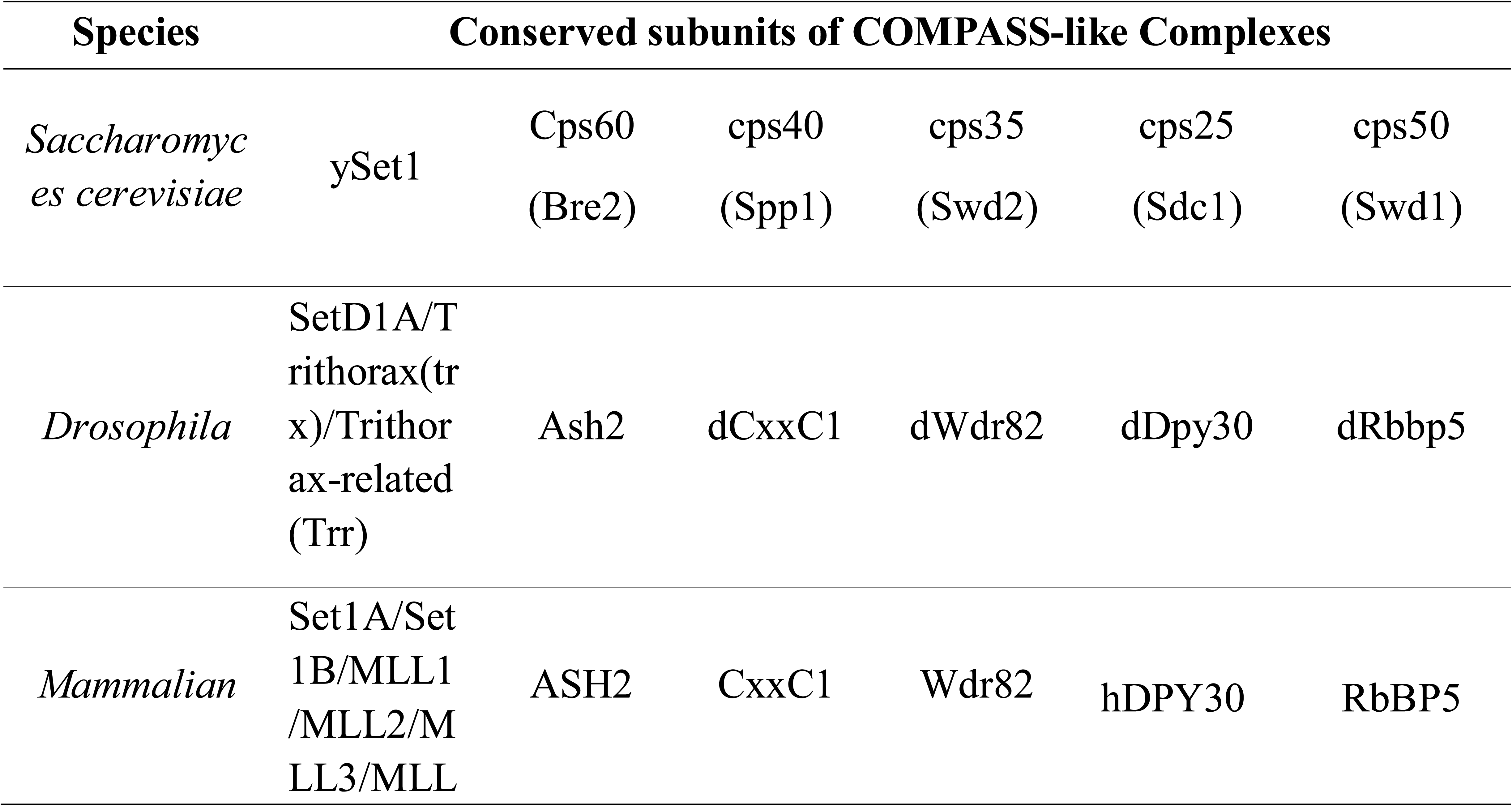
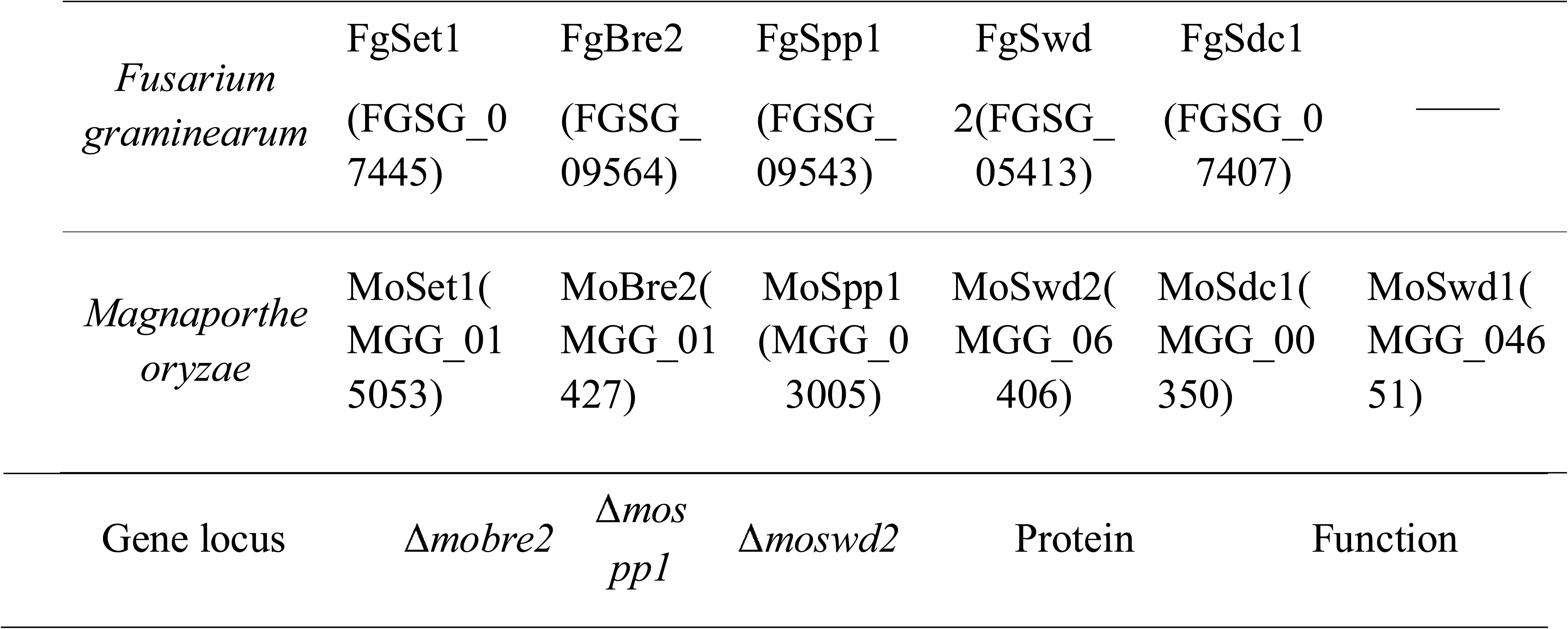
Subunit composition of COMPASS-like family from yeast to human.

## RESULTS

### Phylogenetic analysis and affinity purification of five subunits, MoBre2, MoSwd1, MoSdc1, MoSet1 and MoJmjd2, from the COMPASS-Like complex of *M. oryzae*

To explore whether homologues of the core COMPASS-like complex components are present in *M. oryzae*, we performed a phylogenetic analysis of COMPASS-like homologues from several animals and plants; the results suggested that the subunits of the COMPASS-like complex are highly conserved in *M. oryzae*. We blasted yeast homologues in the *M. oryzae* protein database with the amino acid sequences of proteins such as Cps60/Bre2, Cps50/Swd1, Cps40/Spp1, Cps35/Swd2, Cps25/Sdc, Cps30/WDR5 and Cps30/Swd3. We identified six homologues for these proteins, namely MoSet1 (MGG_15053), MoBre2 (MGG_01427), MoSwd1 (MGG_04651), MoSpp1 (MGG_03005), MoSwd2 (MGG_06406) and MoSdc1 (MGG_00350) (Fig. 1; Table 1). Orthologues of all the components in the COMPASS-like complex are single copies in *M. oryzae*, and their orthologues from representative species, including *S. cerevisiae*, *F. graminearum*, *Arabidopsis*, *Drosophila* and mammalian species (Fig.1; Supporting Information Fig. S1; Table 1; Supporting Information Table S1).

**Fig. 1.**
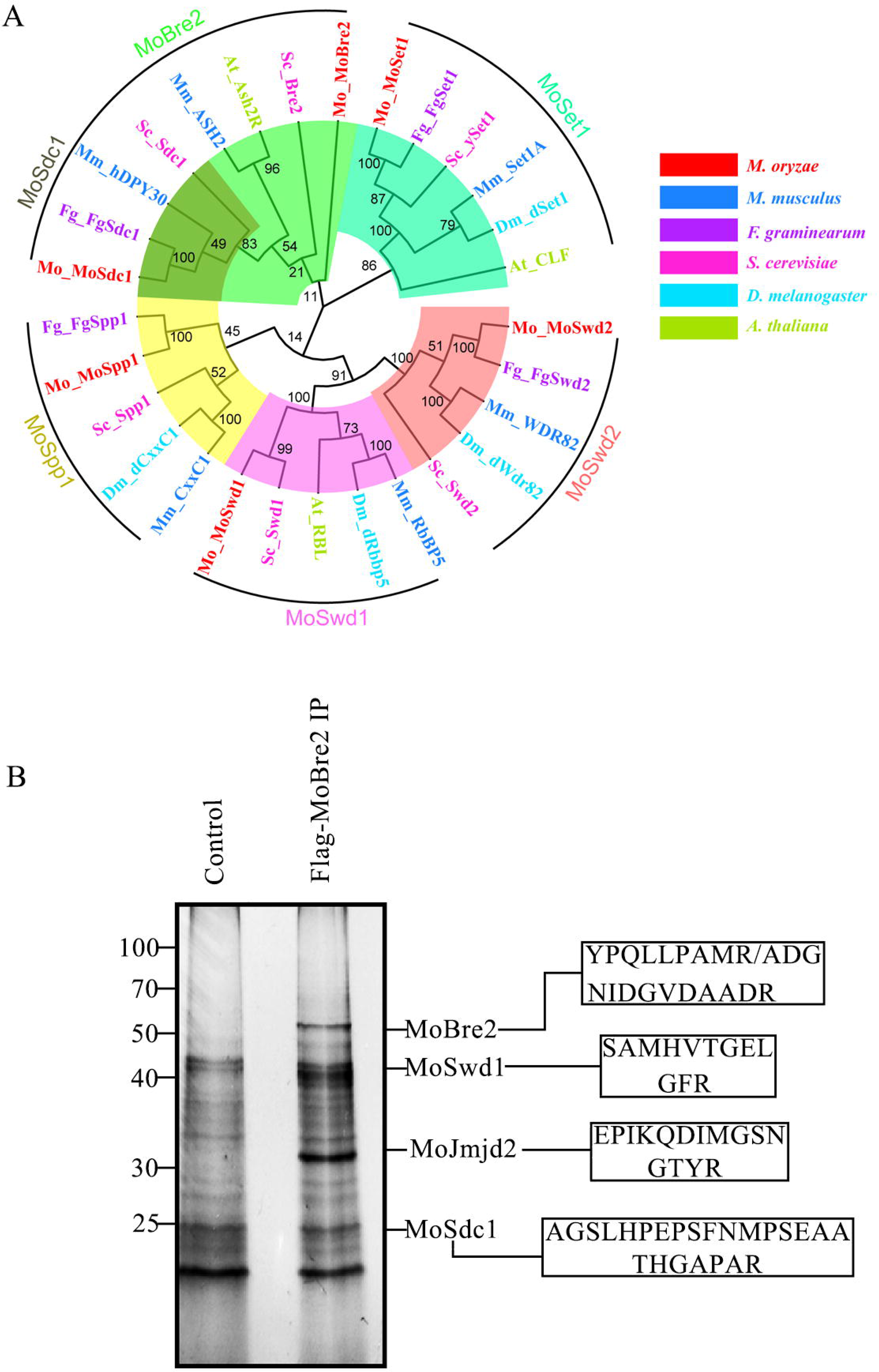
Neighbour-joining tree of candidate COMPASS or COMPASS-like orthologues from *M. oryzae* and other species. Panel A: Co-immunoprecipitation and mass spectrometry analysis of the COMPASS-like complex from *M. oryzae*. Panel B: The boxes contain the peptide sequences identified using tandem mass spectrometry.

To purify the Set1/COMPASS-like complex in *M. oryzae*, we first generated strains able to stably express Flag-MoBre2 and performed an affinity purification using the total nuclear extracts from the Flag-MoBre2 strains (Fig. 1B; Supporting Information Fig. S2). Immunoprecipitated complexes were subjected to silver staining and protein identification using MudPIT (Multidimensional Protein Identification Technology). Tandem mass spectrometry analysis was used to identify the five proteins, MoBre2, MoSwd1, MoSdc1, MoSet1, and a novel JMJC domain-containing protein (MGG_09186) (Supporting Information Table S2). A sequence alignment of MGG_09186 revealed that it contains Jumonji (JmjC) catalytic domains, which can demethylate trimethylated lysine on histones, suggesting that this protein may act as a histone demethylase (Supporting Information Fig. S3A-C). We named MGG_09186 as MoJmjd2 (Smith et al., 2008; Huh et al.,2017). In addition, although we did not detect an obvious MoSet1 band in the SDS-PAGE gels, we did find that our MoBre2-Flag preparations contained significant amounts of MoSet1 (Supporting Information Table S2). However, the MudPIT did not result in peptides unique to other homologous subunits, such as MoSpp1, MoSwd2 and MoSdc1. The results clearly indicate that MoBre2, MoSwd1, MoSdc1, MoSet1 and MoJmjd2 have stable associations with COMPASS-like complexes of *M. oryzae in vivo*.

**Table 2.**
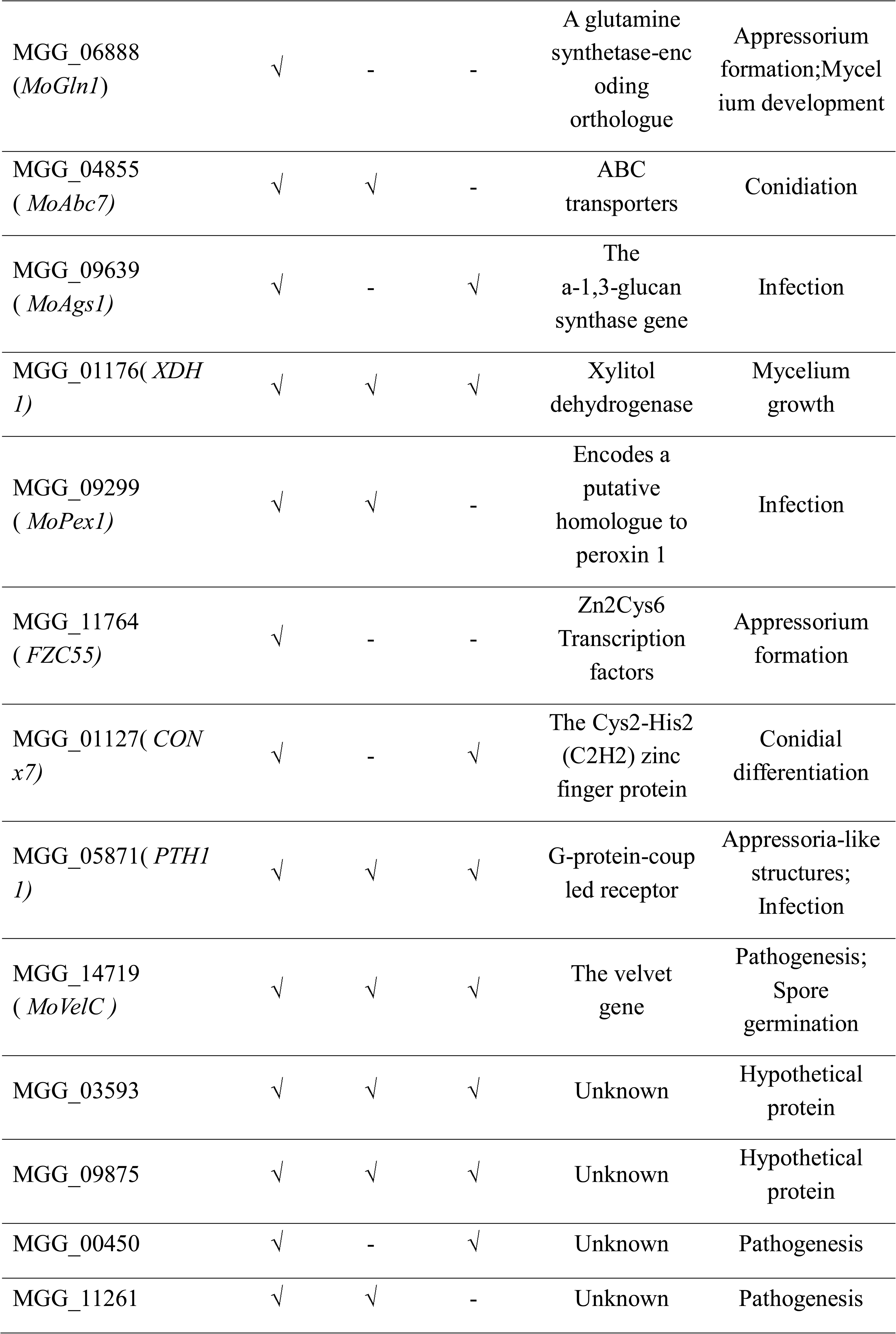
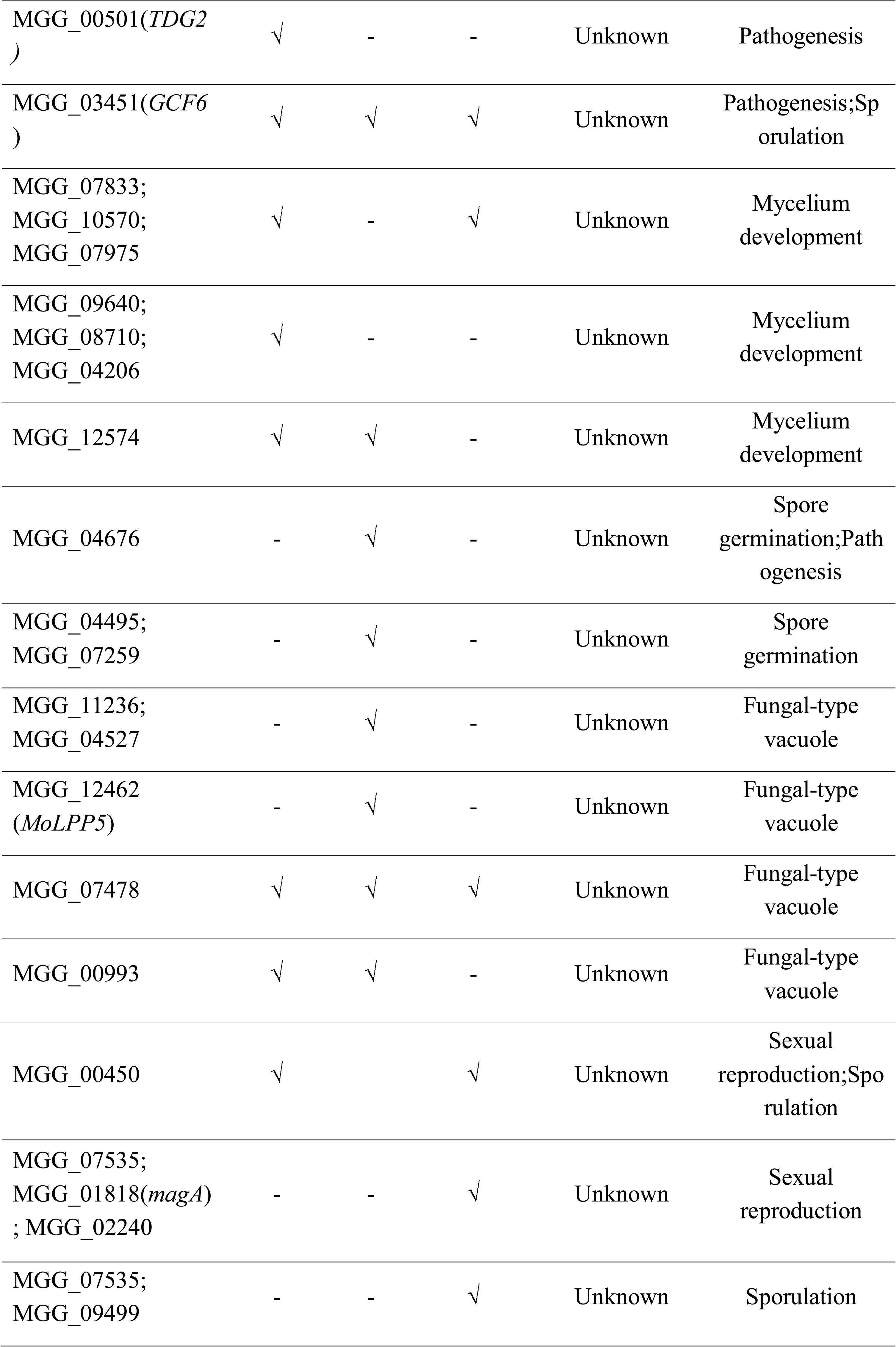
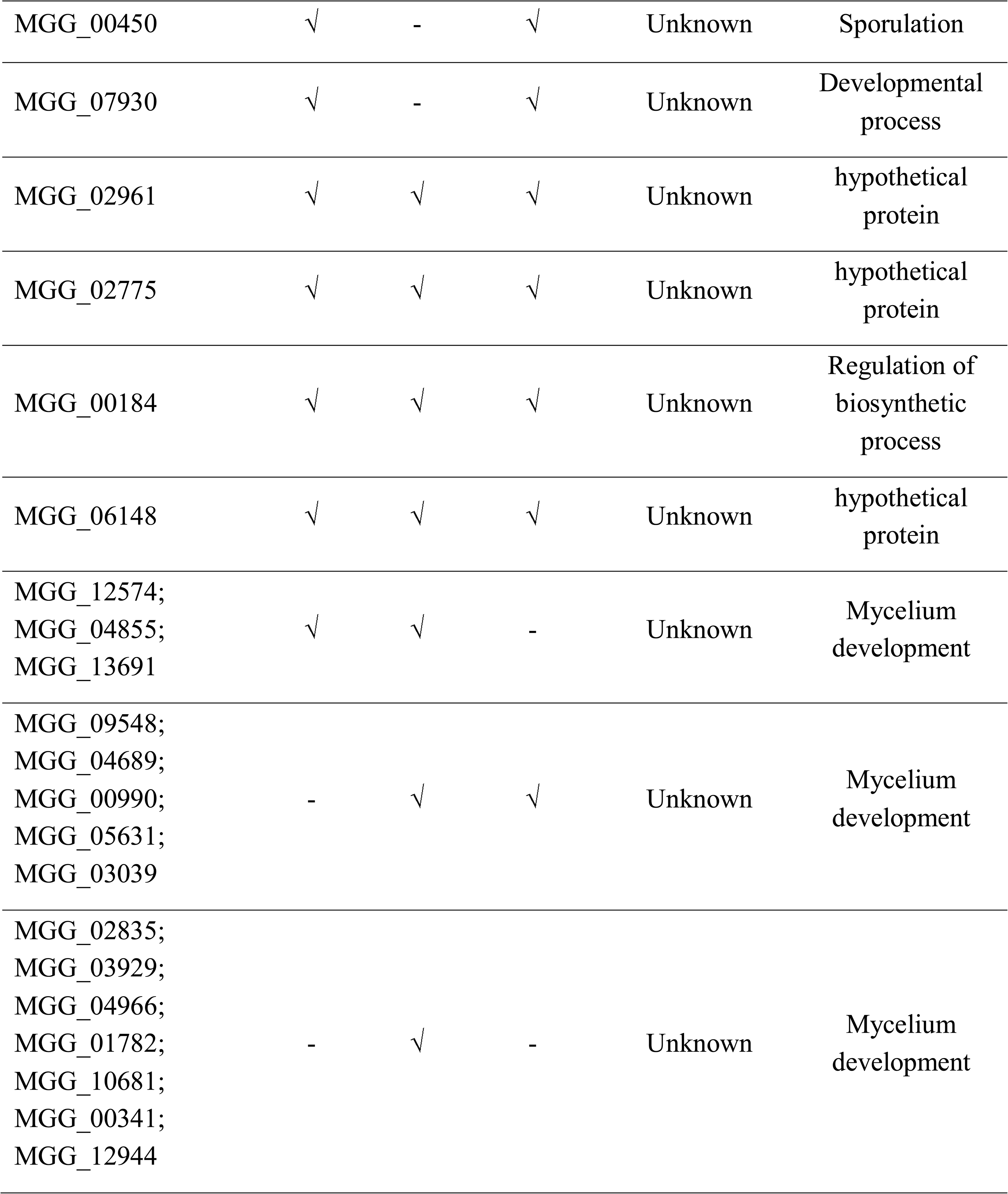
List of the some downstream target genes involved in conidiation, mycelium development, or pathogensis in fungal development and infection in the Δ*mobre2*, Δ*mospp*1 and Δ*moswd2* deletion mutants.

### MoBre2 physically interacts with MoSdc1 via its SPRY domain _401-613_ in vitro and the two subcellularly colocalize *in vivo*

In *M. oryzae*, MoBre2 shares 17.36.% and 12.70% of its identity with yeast Bre2 and human ASH2L, respectively (Supporting Information Fig. S1C-E). In addition, MoSdc1 and yeast Cps25 have a conserved DPY30 domain of approximately 37 amino acids with 68.57% shared identity (Supporting Information Fig. S1D). The MoBre2 protein contains two SPRY domains (Fig. 2A). Recently, the ASH2L-DPY30 interaction was reported to be important for the regulation of H3K4 methylation (Chen et al., 2012; Tremblay et al., 2014; Li et al., 2016).

**Fig. 2.**
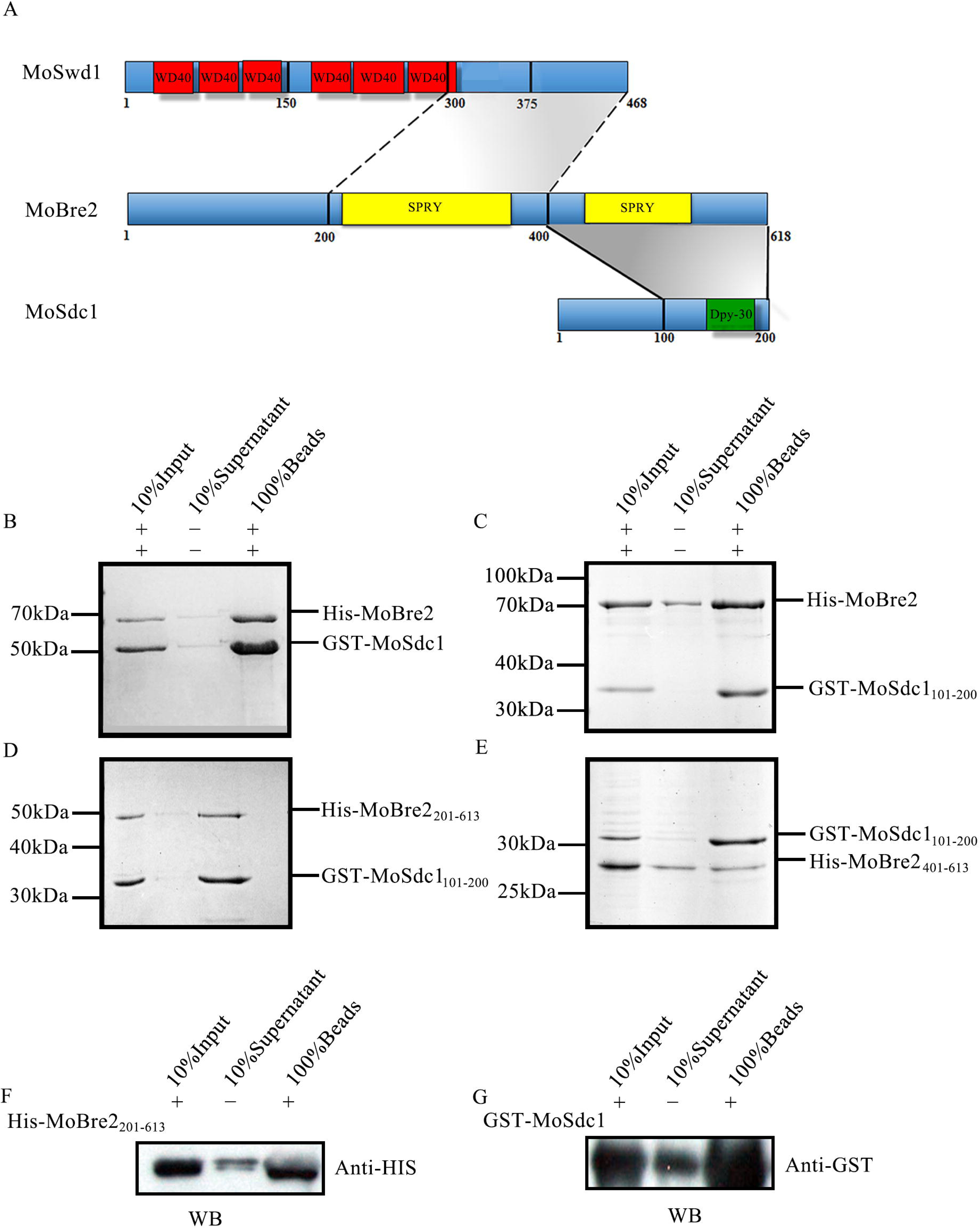
MoBre2 interacts directly with MoSdc1 *in vitro*. Panel A: Domain organization of MoBre2, MoSdc1 and MoSwd1. In MoSwd1, the WD40-repeat domains are shown in red; in MoBre2, the SPRY domains (SPLA and RY anodine receptor domains) are shown in yellow; in MoSdc1, the DPY30 domain is shown in green. Numbers indicate residue numbers at the boundaries of various subdivisions. Protein interactions are indicated with grey areas. Panel B: GST pull-down assay with MoBre2 and MoSdc1. GST-tagged MoSdc1 was incubated with His-tagged MoBre2. The eluates were subjected to SDS-PAGE. Panel C-D: Full-length MoBre2 interacts with the C-terminal region of MoSdc1 (residues 101–200) but not with the N-terminal region of MoSdc1 (residues 1–100). Panel E: The fragments of MoSdc1 (residue 101–200) interact with the two deletion mutants (MoBre2_201-613_, MoBre2_401-613_) Panel F-G: Western blot analysis showing that His-tagged MoBre2*_201-613_* interacts with GST-tagged full-length MoSdc1.

To investigate whether MoBre2 interacts with MoSdc1 or MoSwd1, we expressed and purified recombinant proteins corresponding to the full length, N-terminal and C-terminal regions of MoBre2, MoSdc1 and MoSwd1. Our immunoprecipitation experiments showed that MoBre2 can directly interact with MoSdc1 (Fig. 2A-G; Supporting Information Table S3), but not with MoSwd1 (Supporting Information Fig. S4A, B). To understand which residues are required for interactions between MoBre2 and MoSdc1, we constructed three deletion mutants of MoBre2: (amino acids 1-200, termed MoBre2_1-200_; amino acids 201-613, termed MoBre2_201-613_; and amino acids 401-613, termed MoBre2_401-613_), two deletion mutants of MoSdc1 (amino acids 1-100, termed MoSdc1_1-100_; amino acids 101-200, termed MoSdc1_101-200_) (Fig. 2C-E). As shown in Fig. 2, a pull-down assay showed that MoBre2 retained the binding activity with MoSdc1_101-200_ (Fig. 2C), but not MoSdc1_1-100_ (Supporting Information Fig. S4A).

In addition, MoSdc1_101-200_ physically interacts with MoBre2_201-613_ and MoBre2_401-613_ (Fig. 2D, E), whereas this construct failed to bind to MoBre2_1-200_ (Supporting Information Fig. S4B), which suggests that the SPRY domain located between 401 and 613 of MoBre2 is required for recognition with MoSdc1. Furthermore, a western blot analysis showed that MoSdc1 can interact with MoBre2_401-613_ (Fig. 2F, G). Taken together, these results clearly indicate that a 212-residue SPRY-C-terminal domain of MoBre2 (residues 401-613) is required for interactions with MoSdc1 and that the conserved DPY30-C-terminal regions of MoSdc1 are required for the binding activity with MoBre2 (Fig. 2D, E). However, direct interactions between MoSwd1_1-375_, MoSwd1_150-375_, MoSwd1_300-468_ and MoBre2 was not detected in pulldowns of these protein segments (Supporting Information Fig. S4D-F), suggesting that MoBre2 cannot interact with MoSwd1 directly *in vitro*.

To further explore whether MoBre2 can interact with MoSdc1 *in vivo*, the entire coding regions of MoBre2 and MoSdc1 were fused with green fluorescent protein and red fluorescent protein, respectively. These two constructs were co-transformed into the wild-type strain P131. As shown in Fig. 3, the GFP-MoBre2 was co-localized with the RFP-MoSdc1, clearly showing that MoBre2 can interact with MoSdc1 *in vivo* (Fig. 3). Moreover, these two proteins were localized to the nucleus in the mycelium (Fig. 3A-D), conidia (Fig. 3E-H), appressorium (Fig. 3I-L) and infectious hyphae (Fig. 3M-P). Taken together, these results indicate that MoBre2 can directly interact with MoSdc1 *in vitro* and *in vivo*.

**Fig. 3.**
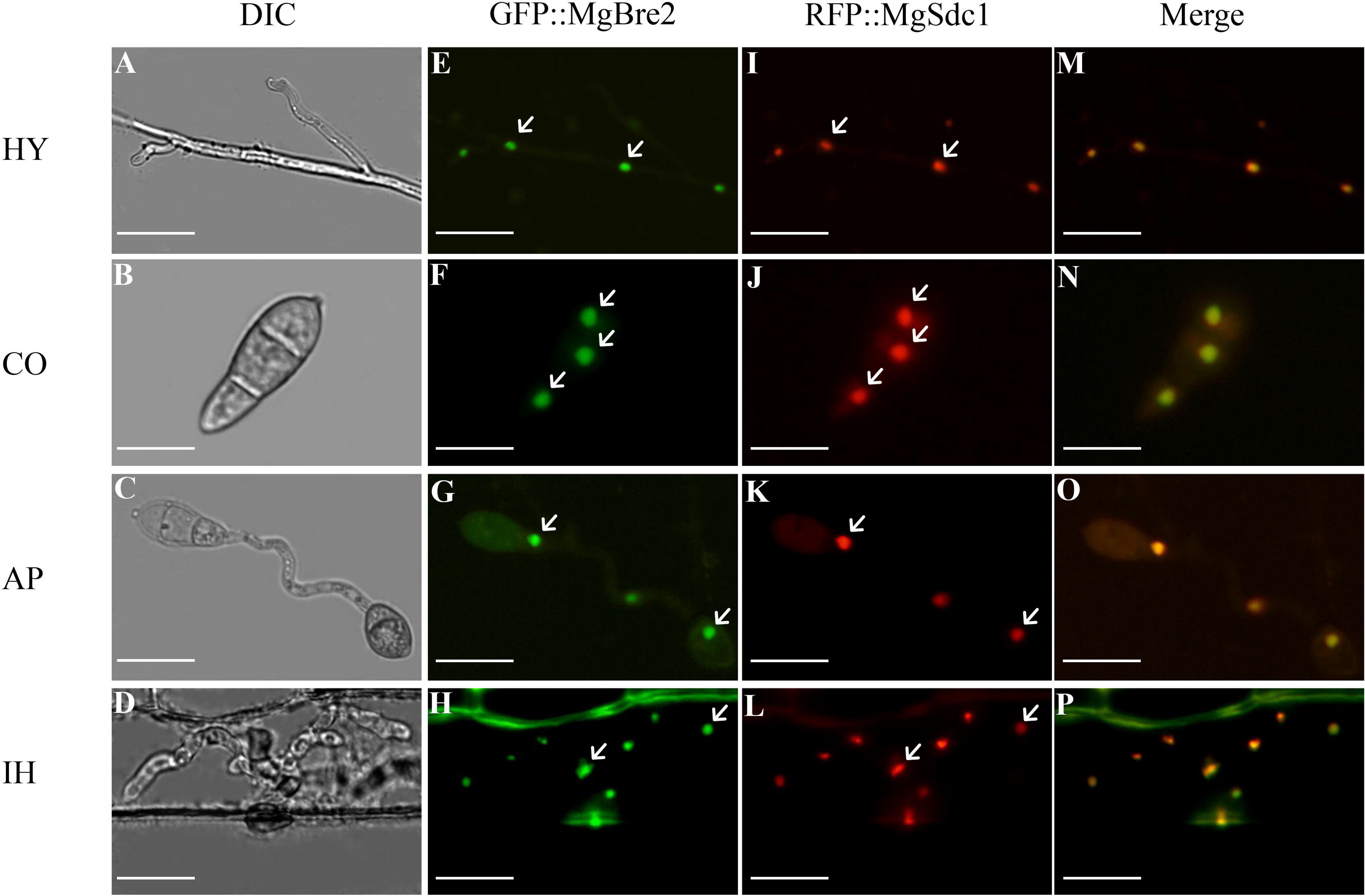
Dual-colour imaging of GFP-MoBre2 and RFP-MoSdc1 in *M. oryzae* by confocal laser scanning microscopy. The GFP-MoBre2 and RFP-MoSdc1 fusion constructs were transformed into the protoplasts of wild-type P131. The GFP-MoBre2 and RFP-MoSdc1 signals were merged. HY, vegetative hyphae; CO, conidia; AP, appressoria; IH, infection hypha. DIC, differential interference contrast. Scale bars = 20 mm.

### MoBre2, MoSwd2 or MoSpp1 are critical for fungal development and invasive hyphae formation during pathogenesis

Next, we used a gene replacement approach to generate null mutants of *M. oryzae* COMPASS-like complex genes. We successfully obtained three null mutant strains for the *MoBre2*, *MoSpp1* and *MoSwd2* genes (Supporting Information Fig. S5). These deletion mutants were confirmed by Southern blots (Supporting Information Fig. S5B, E, I) and PCR analysis (Supporting Information Fig. S5C, F, J). We found that these three gene deletion mutants had defects in colony growth on OTA plates (Fig. 4A). Although the Δ*mobre2* deletion mutant displayed a reduction in colony growth, severe defects were apparent during conidium formation (Fig. 4A, C, D). The Δ*mobre2* mutant produced approximately 30% of the conidia and Δ*moswd2* produced approximately 10% of the conidia compared with the wild-type strain (Fig. 4C). By contrast, the Δ*mospp1* null mutants showed a small reduction in colony growth without conidium formation defects (Fig. 4C).

**Fig. 4.**
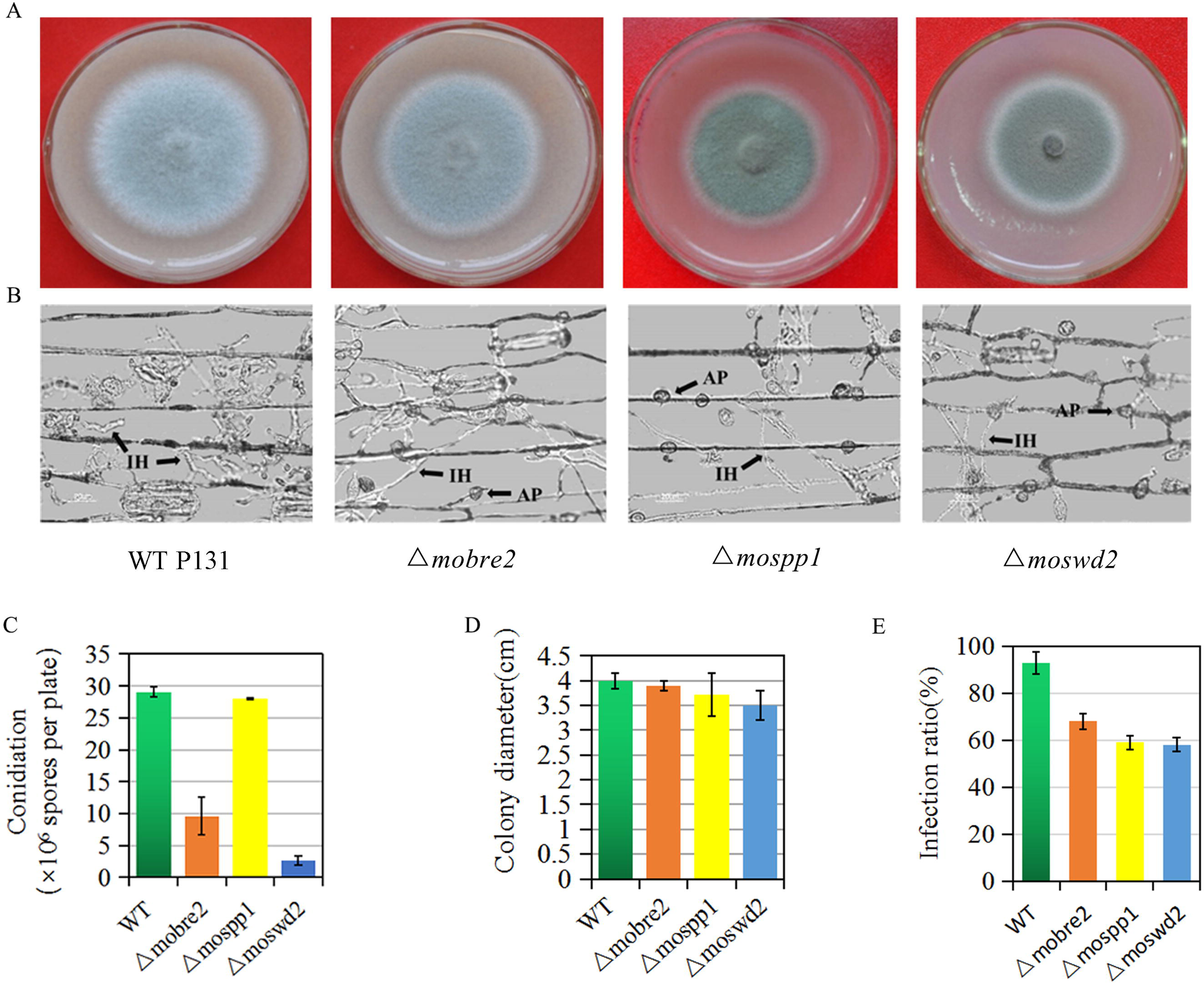
Deletion of *MoBre2*, *MoSpp1* and *MoSwd2* results in defects in hyphal growth, and pathogenic infection. Panel A: Five-day-old cultures of the P131, Δ*mobre2*, Δ*mospp1* and Δ*moswd2* strains grown on oatmeal tomato agar (OTA) medium. Panel B: Barley leaves were sprayed with conidial suspensions (1 × 10^5^ spores/mL) of wild-type P131, the Δ*mobre2*, Δ*mospp1* and Δ*moswd2* deletion mutants 36 h after inoculation. Scale bars = 20 um. Panel C-E: Microscopic observation of conidium formation in the wild-type strain P131 and the Δ*mobre2*, Δ*mospp1* and Δ*moswd2* deletion mutants. Means and SDs were calculated from three independent experiments. *, P < 0.01, n > 100.

To evaluate the virulence of Δ*mobre2*, Δ*moswd2* and Δ*mospp1* null mutants, conidia suspensions of these three null mutants and from wild-type P131 were inoculated onto the surface of barley. These three mutants showed severe defects in their invasive hyphae formation at 36 h after inoculation (Fig. 4B). At 36 h after inoculation, wild-type P131 formed bulbous infection hyphae and could extend to neighbouring host cells on the barley epidermis according to our microscopic observation (Fig. 4B, E). The infection ratio is 68%, 59% or 58.1% in the Δ*mobre2*, Δ*mospp1* and Δ*moswd2* deletion mutants, respectively, compared with wild-type strains (Fig. 4E). However, at 5 d after conidial suspension inoculation onto the seedlings of rice cultivar LTH, the typical robust lesions of rice blast were observed on all three null mutants and wild-type P131 (Supporting Information Fig. S6A, B), which suggests that MoBre2, MoSwd2 and MoSpp1 are required for fungal-host interaction in the early stages of pathogenesis. Taken together, these results indicate that MoBre2, MoSwd2 and MoSpp1 are required for in infection during invasive hyphae formation of the pathogen during fungal development in the early stages of pathogenesis.

### MoBre2, MoSwd2 and MoSpp1 fine-regulate H3K4me3 distribution near TSS target regions

Considering that the COMPASS complex is highly conserved in its H3K4me3 methylation function and activates its target genes expression, we firstly examined the global levels of H3K4me3 in Δ*mobre2*, Δ*mospp1* and Δ*moswd2* gene deletion mutants. A western blot analysis showed that these three null mutants exhibited a robust H3K4me3 decrease throughout the entire genome (Fig. 5, panel A). To determine H3K4me3′s chromatin distribution pattern at the genome-wide level, we performed ChIP-seq (chromatin IP-sequencing) experiments using antibodies against against H3K4me3 in the strains that were devoid of *MoBre2*, *MoSpp1* and *MoSwd2* genes.

**Fig. 5.**
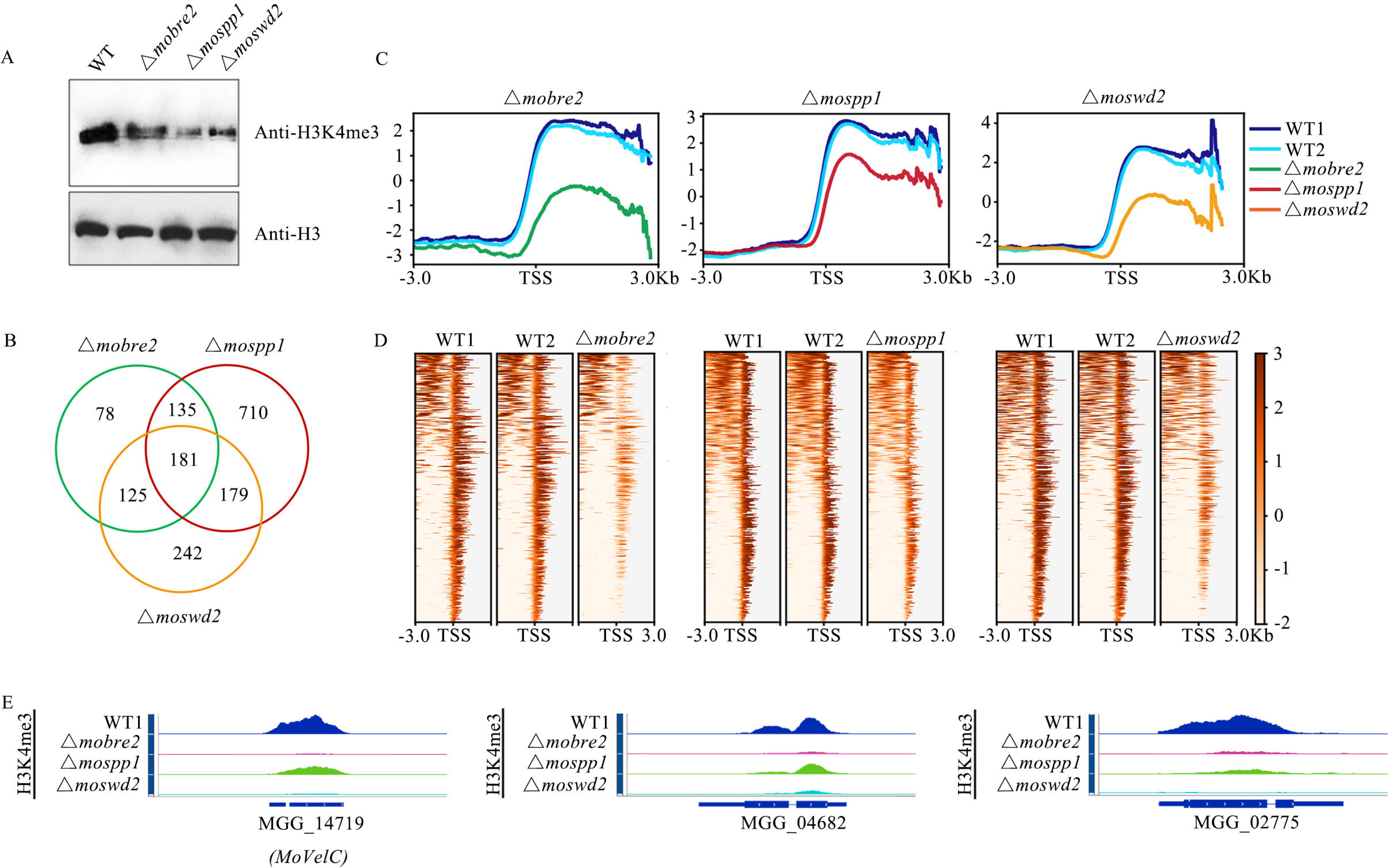
The *MoBre2*, *MoSwd2* and *MoSpp1* deletion mutants display significantly decreased H3K4me3 profiles in TSS target regions. Panel A: Western blot analysis of histone H3K4me3 modifications in the wild-type P131 and Δ*mobre2*, *mospp1* and Δ*moswd2* strains. Panel B: The number in the Venn diagram shows the H3K4me3 decreased peaks in three deletion mutants, compared with the wild-type. Panel C: The H3K4me3-ChIP-seq distribution of enriched peaks around the TSS of the target genes in *MoBre2*, *MoSwd2* and *MoSpp1* deletion mutants. Panel D: The heatmaps show the distribution of H3K4me3-ChIP-seq peak signal density in different samples. Panel E: The read coverage shows that the peaks in these three eletion mutants are significantly decreased compared with the wild-type strain in selected sample genes.

In total, we identified 519, 1205, and 727 H3K4me3-decreased peaks in the Δ*mobre2*, Δ*mospp1* and Δ*moswd2* deletion mutants, respectively, compared with wild-type strains (Fig. 5B; Supporting Information Table S4). Remarkably, close to 60% (316) of the H3K4me3 decreased peaks in the Δ*mobre2* deletion mutant overlapped with the Δ*mospp1* deletion mutant and close to 59% (306) of the H3K4me3 decreased peaks in the Δ*mobre2* deletion mutant overlapped with the Δ*moswd2* deletion mutant (Fig. 5B). Such striking similarity at the genome-wide level strongly suggests a functional connection among the MoBre2, MoSpp1 and MoSwd2.

Significantly, when we sorted all the genes according to their H3K4me3-ChIP-seq reads, we found that the peak distribution was in the TSS regions (−3000 bp to +3000 bp) (Fig. 5C-D). Furthermore, the Δ*mobre2,* Δ*mospp1* and Δ*moswd*2 deletion mutants displayed a highly similar profile in the TSS regions (Fig. 5C-D). The signals of enriched H3K4me3-ChIP-seq reads in the TSS region were largely decreased in deletion mutants compared to the wild-type strain (Fig. 5C). In the Δ*mobre2* mutant, the H3K4me3 signal was reduced 3.5 compared to the wild-type strain (Fig. 5C, left panel), while in the Δ*mospp1* and Δ*moswd2* mutants, the signal displayed 2.6- and 2.2-fold changes respectively (Fig. 5C). These results are further corroborated with decreased intensity of peak signals in heatmaps comparing the knockout mutants with wild-type (Fig. 5D).

We observed that MoBre2, MoSwd2 and MoSpp1 are required for conidiation or infection defects during pathogenesis stage (Fig. 4), and thus we reasoned that MoBre2, MoSwd2 and MoSpp1 might specifically target to fungal development genes. We then performed the comparisons for enriched biological processes in the deletion mutants of Δ*mobre2*, Δ*mospp1* and Δ*moswd2* versus the wild-type strain which showed that many among their target genes are especially involved in mycelium development ((Fig. 5B; Table 2; Supporting Information Fig. S7). In these hypothetical genes, we identified 69/519, 128/1205, 103/727 genes involved in mycelium development and pathogenesis in the *MoBre2*, *MoSpp1* and *MoSwd2* gene deletion mutants, respectively (Supporting Information Table. S5). Among them, there are 64 candidate genes most likely involved in mycelium development were shared by these three deletion mutants (Supporting Information Table S6). Moreover, we selected seven candidate genes from the 181 overlapping genes: *MoGln1* (MGG_06888), *MoAbc7* (MGG_04855), *MoAgs1*(MGG_09639), *XDH1*(MGG_01176), *MoPex1*(MGG_09299), *FZC55*(MGG_11764) and *CONx7*(MGG_01127), all seven are involved in spore germination, fungal growth, or pathogenesis (Fig. 5B; Supporting Information Table S6). MoABC7 has been previously revealed to be involved in virulence, conidiation and abiotic stress tolerance, and MoPEX1 has been shown to be involved in infection-related morphogenesis and pathogenicity. And were previously demonstrated to be involved in mycelium development (Kim *et al*., 2013a; 2013b; Deng et al., 2016; Kou et al., 2016). We also selected three overlapped target genes, MGG_04682, MGG_14719 and MGG_02775, and confirmed that the H3K4me3 coverages are significantly decreased in Δ*mobre2*, Δ*mospp1* and Δ*moswd2* deletion mutants (Fig. 5, panel D). Such a striking similarity of phenotypes among the Δ*mobre2*, Δ*mospp1* and Δ*moswd2* knockouts at the genome level strongly suggests a functional connection among MoBre2, MoSpp1 and MoSwd2.

### Decreased target gene expression caused by deletion of MoBre2, MoSwd2 or MoSpp1 highly correlates with decreased H3K4me3

We further investigated whether the deletion of *MoBre2*, *MoSwd2* or *MoSpp1* would changes in gene expression. We then performed RNA-seq in both wild-type strains and each deletion mutant. Notably, the deletion of *MoBre2* showed relatively moderate transcriptional changes, while the deletion of *MoSpp1* or *MoSwd2* caused more obvious and consistent alterations with the thresholds above 2-fold changes (deletion mutants vs. wild-types) and an False Discovery Ratio (FDR) less than 0.1, including about 42.47% and 55.83% consistently up- and down-regulated genes in Δ*mospp1* or Δ*moswd2* versus wild-type (Fig. 6A; Supporting Information Table S7).

**Fig. 6.**
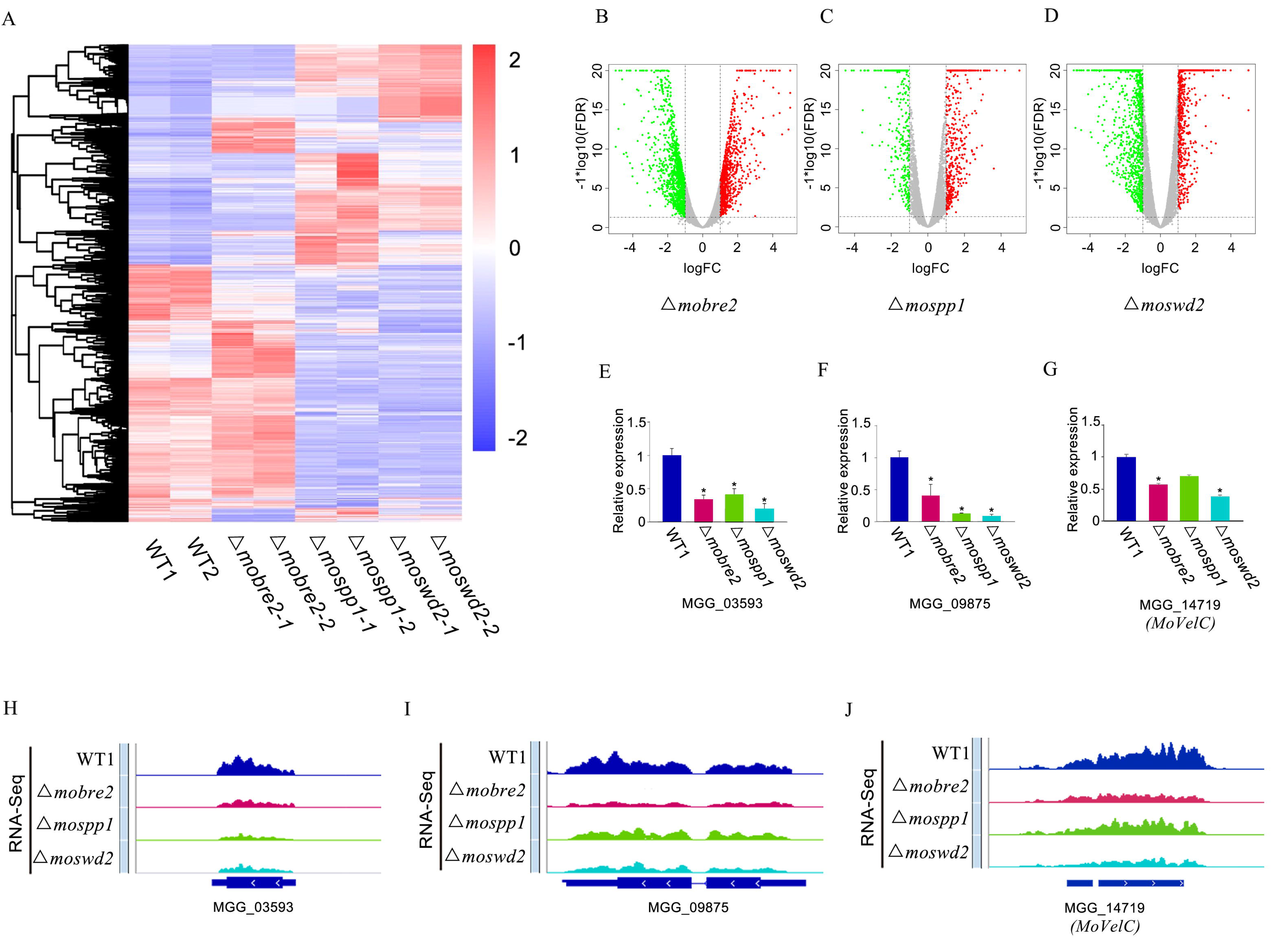
The *MoBre2*, *MoSwd2* or *MoSpp1* gene deletion mutants exhibit gene expression decrease. Panel A: The heat map shows all the differentially expressed genes in the comparison between the wild-type and the three deletion mutants. Panels B-D: The differentially expressed genes are shown in volcano plots, the x-axis shows the fold-changes transfered by log2, and the y-axis shows the Flase Discovery Ratio (FDR) transfered by minus log10. Panels E-G: The read coverage shows that the expression signals of selected targets in these three deletion mutants are significantly decreased compared with the wild-type strain in some selected genes involved in spore germination. Panels E-G: qRT-PCR experiment for spore germination genes show the expression in WT decreased compared with knockouts.

To investigate which biological processes are affected by the *MoBre2*, *MoSpp1* and *MoSwd2* genes, we further performed a gene ontology analysis for the downregulated genes in each knockout strain (Fig. 6B-D; Supporting Information Table S7). Some selected candidate genes such as *PTH11* (MGG_05871), *MoVelC* (MGG_14719) and MGG_03593 are all involved in spore germination and pathogenesis (Supporting Information Table S11; Kim *et al*., 2013a; 2013b; Deng *et al*., 2016; Kou *et al*., 2016). These genes affected in Δ*mobre2*, Δ*mospp1* and Δ*moswd2* were more likely to be involved in developmental processes such as spore germination, transcriptional regulation (transcription factor activities) and pathogenesis (Fig. 6B; Supporting Information Table S8). More importantly, the genes showing decreased expression in the *MoSpp1* or *MoSwd2* deletion mutants are also involved in growth or development of symbionts on or near host, multi-organism processes or other aspects of pathogenesis (Fig. 6B-D). The functions of these selected genes are clearly consistent with the phenotypes we have observed in Δ*mobre2*, Δ*mospp1* or Δ*moswd2* deletion mutants, that have defects in colony growth, infection, and other related aspects of early pathogenesis (Fig. 4B).

We detailed gene expression patterns of three genes, MGG_03593, *MoVelC* (MGG_14719) and MGG_09875. The read coverage of these genes indicated a significant drop in expression compared with the wild-type strain (Fig. 6E-G). Furthermore, qRT-PCR experiments for these genes selected indicate that expression in the wild-type was higher than in the knockouts (Fig. 6H-J).

To further observe the relationship between gene expression changes and H3K4me3 changes, we overlapped the downregulated genes in Δ*mobre2*, Δ*mospp1* and Δ*moswd2* deletion mutants. For each comparison between the deletion mutant and the wild-type (Fig. 7A), we observed that the downregulated genes were likely to have decreased the H3K4me3 peaks, which suggests that the downregulated genes that are overlapped targets of the MoBre2, MoSpp1 and MoSwd2. In Δ*mobre2* deletion mutants, there are 8 (3.5%) and 12 (5.3%) genes are overlapped with the Δ*mospp1* and Δ*moswd2* deletion mutants, respectively (Fig. 7, panel A). Notably, five overlapped genes that had decreased expression, MGG_02961, MGG_02775, MGG_00184, *MoVelC* (MGG_14719) and MGG_06148, followed the same trend in each of the Δ*mobre2*, Δ*mospp1* and Δ*moswd2* knockout lines (Fig. 7A, B; Supporting Information Table S9). We identified expression levels through RNA-seq for those genes with decreased H3K4me3 peaks and found that these genes are also consistently down-regulated in the knockout strains (Fig. 7B). We further confirmed that a gene, we obtained (Fig. 7A), with reduced gene expression in RNA-seq had a similar H3K4me3 reduction pattern in ChIP-seq (Fig. 7A, C). For MGG_02775, *MoVelC* (MGG_14719) and MGG_04682, the read coverage of these genes indicated a significant drop both in gene expression and H3K4me3 occupancy (Fig. 7D, E). Significantly, the majority of H3K4me3-decreased peak target genes had significant expression changes, which correlates with the well-known effects of histone modification on gene expression.

**Fig. 7.**
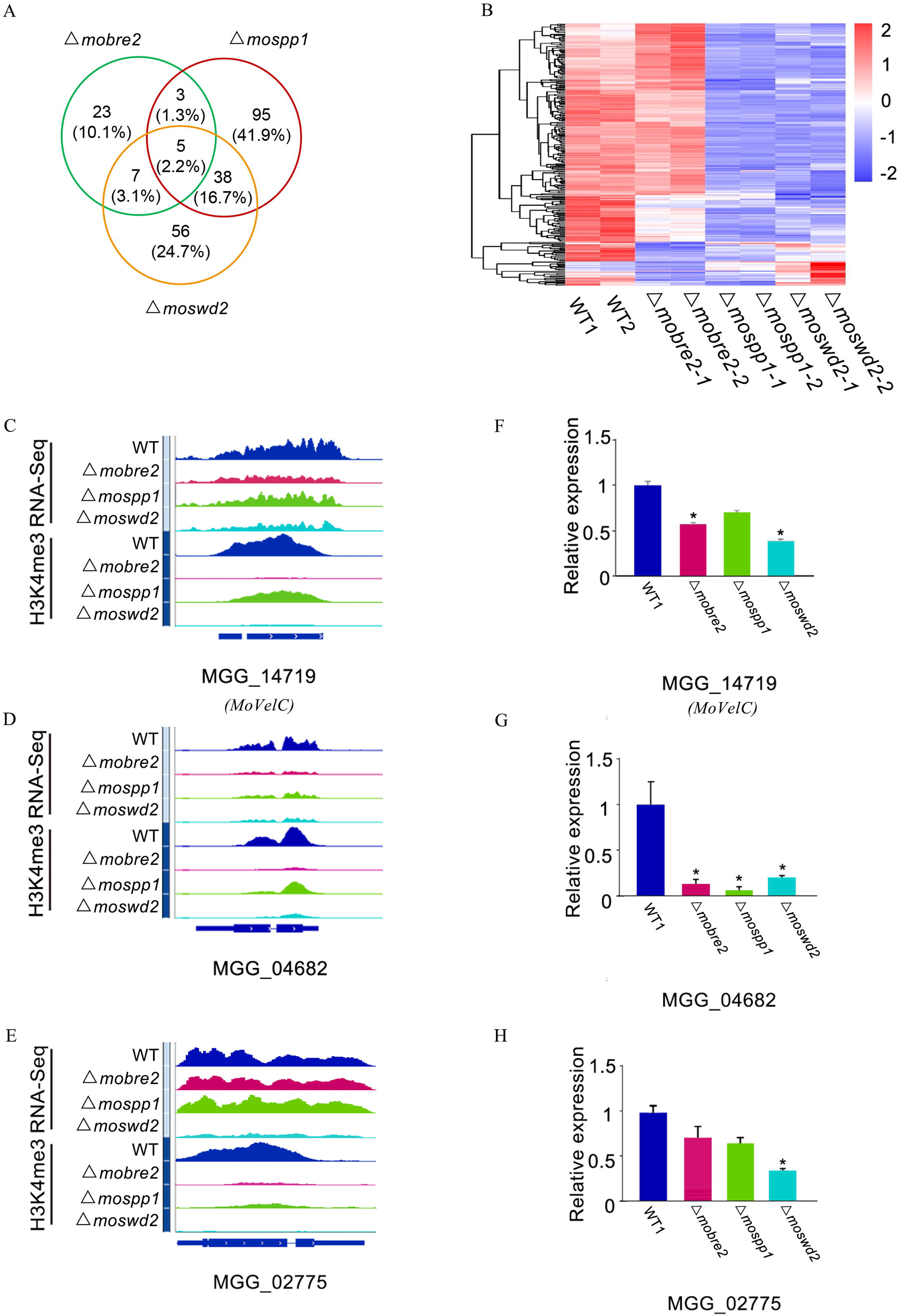
Decreased gene expression in the MoBre2, MoSwd2 or MoSpp1 gene deletion mutants is highly correlated with decreased H3K4me3. Panel A: Venn diagrams show changes in down-regulated gene expression in knockouts compared to the wild-type strains. The numbers of down regulated genes are indicated. Panel B: Heat map of differentially expressed genes between knockouts and the wild-type strains. Panels C-E: Examples of genomic tracks showing genes with decreased H3K4me3-ChIP-seq signal levels and downregulated RNA-seq reads. Panels F-H: qRT-PCR experiment for selected genes show the expression in WT decreased compared with deletion mutants. Identified genes show decreased signals and the expression level was reduced in deletion mutants.

Together, these results suggest that the MoBre2, MoSpp1 and MoSwd2 may assemble into a functional COMPASS complex and mediate H3K4me3-regulated gene expression.

## DISCUSSION

### Highly conserved roles of COMPASS-Like H3K4 methyltransferase complexes in *M. oryzae*

This study is the first to uncover the molecular mechanism of the COMPASS-like complex in a plant pathogenic fungus providing new insight into the epigenetic regulatory mechanism of target gene expression during fungal development and infection-related pathogenesis.

In fungi, *F. graminearum* has been found to contain four COMPASS-like genes (Jiang et al., 2011; Mohan et al., 2011; Kanda et al., 2013; Liu et al., 2015). Our biochemical studies have demonstrated that the *M. oryzae* complexes are very similar to their mammalian counterparts in terms of subunit composition. We found that all the COMPASS family members are also highly conserved in *M. oryzae*. There are several common subunits, namely MoBre2, MoSwd1, MoSdc1, MoSet1, MoSpp1 and MoSwd2, that are homologues of Cps60, Cps50, Cps25, ySet1, Cps40 and Cps35, respectively (Fig. 1; Table 1). A study showed that MoSet1, a component of COMPASS, is required for global gene expression during infection-related morphogenesis, which is consistent with the findings presented here (Pham et al., 2015). However, we were unable to identify the homologues of yeast Cps30/Wdr5 in *M. oryzae.* Moreover, we did not purify any Cps30-like protein from the immunoprecipitation assay. Further study is required to determine whether there are some Cps30-like proteins in the *M. oryzae* genome.

Previously published crystal structures of COMPASS or COMPASS-like in yeast and humans revealed that a Y-shaped architecture with Cps50 and Cps30 localizes on the top of two adjacent lobes and Cps60-Cps25, forming the base at the bottom (Takahashi et al., 2011; Zhang et al., 2014). In *M. oryzae*, our results also showed that MoBre2/Cps60 can interact directly with MoSdc1 *in vitro*, but not with MoSwd2. Although we did not find that MoSwd1 can interact with MoBre2 in a pull down assay, we did find that Flag-MoBre2 can be co-purified with MoSwd1 in an immunoprecipitation assay *in vivo* (Supporting Information Table S4), which suggests that MoSwd1 does indeed function in the COMPASS complex of *M. oryzae*.

Interestingly, we also found that our Flag-MoBre2 preparations contained significant MoJmjd2 enrichment (Fig. 1; Supporting Information Table S5). A BLAST protein search revealed a significant similarity between MoJmjd2 and UTX/UTY. Recently, some studies have showed that histone demethylases, such as KdmA, an H3K9/36me3 demethylase, play a role in transcriptional regulation in *A. nidulans* (Agger et a*l*., 2007 & 2019; Matthews et al., 2015). This finding suggests that these UTX-like containing COMPASS-like complexes and their associated bivalent H3K9/27 demethylase and H3K4me3 methyltransferase activities may be involved in dynamic target gene activation (Bernstein et al., 2006; Amar et al., 2015;Bachleitner et al., 2019).

### Contribution by COMPASS-Like complex components provides novel insights into the regulation of target genes in TSS regions

In this study, we found that the MoBre2, MoSpp1, and MoSwd2 are essential for colony growth, conidiation, and invasive hyphae formation in *M. oryzae* (Fig. 6A-E). The three gene deletion mutants exhibited pleiotropic defects as described for the Δ*moset1* mutants (Pham et al., 2015). According to the ChIP-seq and RNA-seq results, several previously characterized target genes, such as *MoABC7*, *MoPEX1*, *FZC55*, *MoAGS1*, and *GLN1*, were among the COMPASS-like complex-downregulated genes with H3K4me3-decreased peaks. In Δ*moswd2* strains, the expression of some target genes involved in conidiation and pathogenesis is decreased, and this finding is consistent with the mutant defects in invasive hyphae formation (Fig. 6B, E; Supporting Information Table S8).

More interestingly, we also found downregulated expression of *MgHTF*1 (MGG_00184), a gene known to regulate conidial production. The Δ*MgHTF*1 mutant failed to form conidia and lost pathogenicity (Liu et al., 2010). The expression of *MgCONx*2 (MGG_02775), a gene that encodes a zinc finger C2H2-like protein, also decreased. The *MgCONx2* mutant was previously deemed essential for conidiogenesis and pathogenesis in *M. oryzae* (Zhang et al., 2014) (Fig. 7; Supporting Information Table S8). The ChIP-seq results show that *MgHTF*1 and *MgCONx*2 both have H3K4me3-decreased peaks in the Δ*moswd2* strains, which suggests that MoSwd2 activates H3K4me3 in the TSS regions to regulate the transcription of target genes (Figs. 6 and 7; Supporting Information Table S4). The expression levels of *MgAnd*1 (MGG_02961) and *MgFox*2 (MGG_06148) were also significantly downregulated in the Δ*mospp1* mutant. *MgAnd*1 encodes a cell cortex protein involved in nuclear migration and positioning. Deleting of *MgAnd*1 from *M. oryzae* leads to a delay in the penetration of the host plants (Rahman et al., 2014). *MgFox*2 encodes a multifunctional enzyme that catalyses the second and third enzymatic reactions of the β-oxidation pathway, and its disruption also leads to a significant reduction in virulence (Patkar et al., 2012).

Notably, among the genes affected by the COMPASS-like complex, the gene *MgVelC* (MGG_14719) was reported to be a positive regulator of conidiation and pathogenicity in *M. oryzae* (Kim et al., 2014). The expression of *MgVelC* was downregulated in both Δ*mobre2 and* Δ*moswd2* gene deletion strains. Furthermore, the ChIP-seq results also show that *MgVelC* has H3K4me3-negative peaks in both gene deletion mutants. These results suggest that *MgVelC* is a potential target gene that is regulated by the COMPASS-like complex. The transcriptional deficiency of these genes may be the cause of severe defects in terms of colony growth, conidiation, and pathogenicity in different mutants (Dang et al., 2001; Wang et al., 2012; D’Urso et al., 2016). Collectively, these findings suggest that the COMPASS-like complex controls a good number of genes for fungal pathogenicity and development in *M. oryzae*. However, the details of how these target genes regulate pathogenesis remain unknown.

In conclusion, our biological, biochemical, and proteomic studies identified novel components of the COMPASS-like complex and elaborated on its contribution to H3K4me3-mediated target gene expression in *M. oryzae*. This work serves as a functional resource for studying infection processes in plant pathogenic fungi and provides a greater breadth of knowledge in epigenetics regarding gene regulatory mechanisms.

## EXPERIMENTAL PROCEDURES

### Strains and culture conditions

The *M. oryzae* wild-type strain P131 was used in this study. The wild-type strain and corresponding transformants were grown on oatmeal tomato agar (OTA) medium and cultured at 25°C under light conditions. For extraction of genomic DNA, RNA, and protein, as well as the isolation of protoplasts, fresh mycelia were broken and cultured in liquid CM media (0.6% yeast extract, 0.3% enzymatic casein hydrolysate, 0.3% acidic casein hydrolysate, and 1% glucose) and shaken at 150 rpm at 25°C for 36 h. For measuring colony sizes, mycelial plugs of 5 mm were inoculated into the center of OTA plates. Colony diameter on each plate was measured for three days. Each experiment was repeated three times.

### Conidiation experiments and virulence tests of M. oryzae

The wild-type, deletion mutant and complementation strains were cultured on OTA plates at 25 °C for four days. Then, 2 mL ddH2O was added to each plate and mycelia were collected with an inoculation loop and transferred to a new OTA plate and dried. The plates were covered with three layers of gauze and cultured at 25°C for 12–16 h. Two mL of ddH_2_O was added to each dish and conidia were scraped with sterile cotton swabs. Conidia suspensions were transferred into a new 50 mL tube and centrifuged for 5 min at a minimum of 5,000 g at 25°C. The supernatant was removed and pellets were resuspended to obtain 2 × 10^4^ conidia per mL in 0.025% (v/v) Tween-20 solution. A blood cell count board was used to count conidia under the optical microscope. For the infection assay, four-week-old seedlings of rice (*Oryza sativa* cv LTH) and eight-day-old seedlings of barley (*Hordeum vulgare*) cv. E9 were used for spray infection assays. Plant incubation was performed as described previously (Cao et al., 2016). All inoculation and conidiation experiments were repeated three times independently.

### Western blot analysis

For all tested strains, total proteins were extracted as described previously (Cao et al., 2016), then separated by 13% denaturing polyacrylamide gel (SDS-PAGE), transferred to an Immobilon-P transfer membrane (Millipore, Billerica, MA, USA) with a Bio-Rad electroblotting apparatus, and evaluated using the antibodies anti-Histone H3 (Abcam, ab1791) and anti-H3K4me3 (Abcam, ab8580), respectively. Samples were incubated with a secondary antibody (Santa Cruz, sc-2313) for chemiluminescent detection.

### Co-IP and silver staining

Immunoprecipitation experiments were performed using a 3 × *MoBre2-Flag* transgenic strain. About 0.5 g seven-day-old mycelia from the FLAG-MoBre2-expressing strain were harvested and ground in liquid nitrogen. Subsequently, total proteins were extracted with protein extraction buffer (50 mM Tris-HCl [pH=7.5], 100 mM NaCl, 5 mM EDTA, 1% Triton-X100, 2 mM PMSF). Then, 15 mL protein extracts were incubated in a 50 μL slurry of anti-FLAG M2 affinity gel (Sigma, A2220) and rotated at 4°C overnight. The beads were washed by 150 mL BC100 (20 mM Tris-HCl [pH=8.0], 100 mM NaCl, 1 mM EDTA, 0.1% NP-40)(Wang et al., 2011). Subsequently, target proteins were eluted with the FLAG peptide (DYKDDDDK). The samples were then boiled in the SDS-PAGE loading buffer and separated in an SDS-PAGE gel. Silver staining experiments were performed using the ProteoSilver plus silver stain kit (Sigma, PROT-SIL2). The silver stained bands were analyzed by mass spectrometry.

### Protein expression in E. coli and purification

The recombinant deletion vectors of MoBre2, MoSdc1 and MoSwd2 were transformed into Rosetta (DE3) *E. coli* bacteria (Biomed, BC204-02, China). After sequencing to determine positive clones, the bacteria were incubated at 37°C at 220 rpm to OD_600_ = 0.6 with Luria-Bertani (LB) medium, then induced by the addition of 1 mM β-D-1-thiogalactopyranoside (IPTG) and incubated at 16°C for 14 h to express GST-tagged fusion proteins. The fusion proteins were purified using Glutathione-Sepharose 4B (GE Healthcare, 17-0756-01) following the manufacturer’s instructions.

### In vitro pull-down assay

The fusion proteins were purified as described previously (Mao et al., 2014). Then, equivalent protein of His-*MoBre2*_FL_ and GST-*MoSdc1*_FL_ fusion proteins were added to a 1.5 mL centrifuge tube and the solution was brought to 400 μL by BC100 (20 mM Tris-HCl [pH=8.0], 100 mM NaCl, 1 mM EDTA, 0.1% NP-40). Next, 40 μL Glutathione-Sepharose 4B (GE Healthcare, 17-0756-01) was added to the solution. The tubes were then rotated and incubated for 4 h at 4 °C to pull down GST-tagged fusion proteins. This was followed by four washes with BC500 (20 mM Tris-HCl [pH=8.0], 500 mM NaCl, 1 mM EDTA, 0.1% NP-40). The beads were boiled in SDS loading buffer for 10 min, centrifuged at 3500 rpm at 4 °C for 5 min, and then the supernatant was separated by 13% denaturing polyacrylamide gel (SDS-PAGE). Finally, the gel was stained with Coomassie Brilliant Blue R-250 (AMRESCO, 0472-10G) and Western blotting using anti-His (Transgen Biotech, HT501) and anti-GST antibodies (Transgen Biotech, HT601) for observation.

### Subcellular localization analysis

For colocalization analysis of MoBre2 and MoSdc1, the full-length of *MoBre2* was amplified by PCR from wild-type P131 cDNA with primer pairs *MoBre2-*GFP-F and *MoBre2-*GFP-R. The resulting fragment and pKNTGRP27 vector were digested with *Eco*RI and *Bam*HI and *MoBre2* was cloned into pKNTGRP27, which was generated by cloning the strong, constitutive RP27 promoter into pKNTG. The full-length of *MoSdc1* was amplified by PCR from wild-type P131 cDNA with primer pairs MoSdc1-RFP-F and MoSdc1-RFP-R. The resulting fragment and pKS vector were digested with *Eco*RI and *Bam*HI, *MoSdc1* was cloned into pKS and the RFP-MoSdc1 fusion construct was generated. The GFP-MoBre2 and RFP-MoSdc1 fusion constructs were digested with *Not* I and co-transformed into wild-type P131. Colocalization of GFP-MoBre2 and RFP-MoSdc1 fusion proteins was performed with a Nikon A1 laser scanning confocal microscope.

### ChIP-seq

Protoplasts for ChIP sequencing were prepared as described previously (Cao et al., 2016). Briefly, the protoplasts were crosslinked with 37% formaldehyde for 10 min, followed by termination of the crosslinking reaction with 10 × glycine for 5 min. Protoplasts were collected by centrifugation at 4000 rpm for 15 min, washed with 0.7 M NaCl, and diluted to 1 × 10^8^ /mL. Centrifugation and suspension of protoplasts occurred with 750 μL RIPA buffer (50 mM Tris-HCl [pH=8.0], 150 mM NaCl, 2 mM EDTA, 1% NP-40, 0.5% sodium dexycholate, 0.1% SDS). The chromatin was sheared by sonication with JY 92-IIDN (SCIENTZJY, China) for 8 min (25% W, output 3S, Stop 5S). Chromatin immunopurification with anti-H3K4me3 antibody (abcam ab8580) and Dynabeads Protein A (Invitrogen 10002D) were incubated overnight at 4 °C as previously described (Mao *et al*., 2014). Subsequently, DNA fragments were extracted using the phenol-chloroform method for constructing an Illumina sequencing library and sequenced with single-ends on a Hiseq 2000.

### Mass spectrometry (LC-MS/MS) and data analysis

Samples were destained in Proteo Silver Destainer A and Proteo Silver Destainer B, and then rinsed with pipetted water several times. Proteins were digested with 500 ng of sequencing-grade modified trypsin (Promega), and the resulting peptides were resolved on a nano-capillary reverse phase column. Using a water: acetonitrile gradient system, at 300 nL/min, sample was directly introduced into an ion-trap mass spectrometer in the data-dependent mode to collect MS/MS spectra on the five most intense ions from each full MS scan. MS/MS spectra were matched to peptides in a database containing *M. oryzae* protein sequences. All proteins with a Protein Prophet probability score of 0.9 were considered positive identifications.

### RNA-seq

Total RNA was isolated from the tissue of wild-type strain P131, Δ*Mobre2*, Δ*mospp1* and Δ*moswd2* deletion mutants, and mRNA was then isolated using a Poly (A) Purist MAG kit (Ambion). Each sample was analysed in technical triplicates, and three experiments were replicated. DNA was removed by RNase-free DNase (Qiagen) followed by column clean-up according to the manufacturer’s instructions. Illumina TruSeq RNA Sample Preparation kits were used to construct RNA-seq libraries; cDNA was sequenced on an Illumina Hiseq 2000.

### Bioinformatics

ChIP-seq read quality was assessed using FASTQC (0.11.7) and aligned to *M. oryzae* genome assembly (MG8) using Bowtie2 software (2.3.4.3). Only uniquely mapped reads were kept for analysis. H3K4me3 peaks were called by MACS2 (2.1.1.20160309) with read extension to average fragment size and the threshold of FDR<=1E-6. Based on the MACS2-called peaks and aligned read files, DiffBind (v2.6.6) was used to identify differential binding peaks of H3K4me3 with FDR<0.01 and binding fold change ≥ 2.

RNA-seq reads were aligned to *M. oryzae* transcriptome annotation (MG8) with the STAR aligner (2.6.0c). Differentially expressed genes and quantification were done using the edgeR package (3.20.9). Plots and charts were drawn with R software and MS Excel.

### Sequence alignment and structural modeling

The sequences of the Unknown and JMJD2A protein sequences were aligned in ClustalX2 and then coloured by ESPript 3.0 (http://espript.ibcp.fr/ESPript/ESPript/; Robert et al., 2014). Using the online phyre2 modelling machine (http://www.sbg.bio.ic.ac.uk/phyre2/html/ page.cgiid=index; Kelley et al., 2015), we constructed and selected the complex of the core Tudor domain of Unknown and H3K27me3 with the highest score, based on the template of the JMJD2A Tudor domain–H3K4me3 complex (PDB code: 2006; Ng et al., 2007).

## Acknowledgements

This work was supported by National Natural Science Foundation of China (Grant no. 31871638), the Special Scientific Research Project of Beijing Agriculture University (YQ201603), the Scientific Project of Beijing Educational Committee (KM201610020005), the Research Fund for Academic Degree & Graduate Education of Beijing University of Agriculture (2019YJS037), the National Laboratory of Biomacromolecules, Institute of Biophysics, Chinese Academy of Sciences, and the National Natural Science Foundation of China (Grant no. 31872916).

## Author contributions

W.X.W designed the experiments. S.D.Z., W.Y.S., M.Y.Z., S.P., X.Y.Z.,Y.Y., D.H., M.S., J.Y., L.E.C., and Y.X. performed the biological experiments. X.Y.L. and Q.Z. performed the bioinformatics analysis. W.X.W wrote the manuscript.

## Supporting Information

**Fig. S1.**
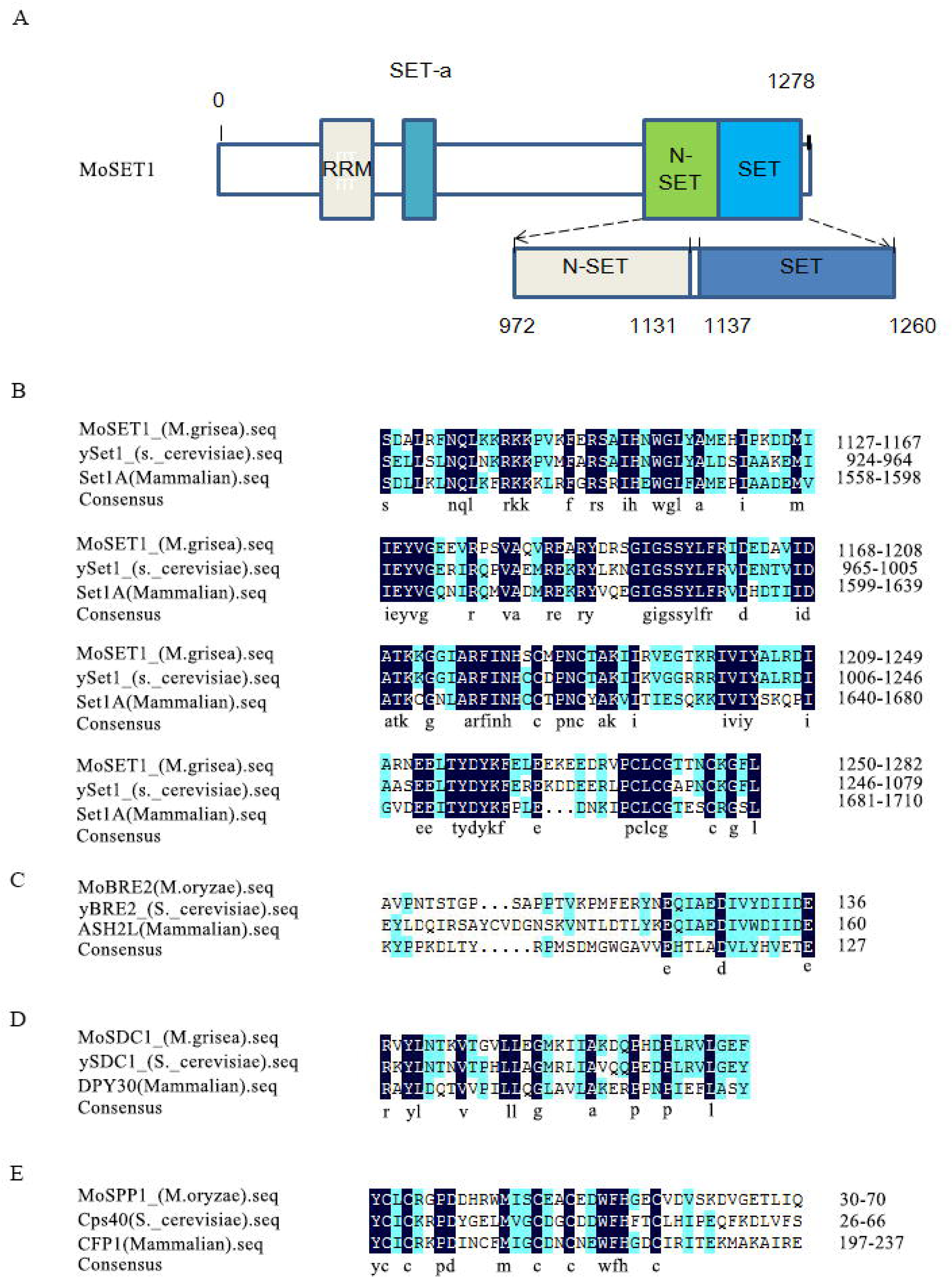
Identification of the H3K4 methyltransferase MoSet1, MoBre2, MoSdc1 and MoSpp1 in *M. oryzae*. Panel A. Schematic representation of MoSet1. Each functional domain is highlighted differently. RRM, RNA recognition motif, N-SET, N-Set domain, SET, SET domain. Panel B. The MoSet1 SET domain is evolutionarily conserved. Sequence alignment of the SET domains from mammals (Set1A), *Saccharomyces cerevisiae* (ySet1) and *M. oryzae* (MoSet1). Positions with 100% and 99-75% amino acid conservation are represented in black and blue, respectively. Panel C. Alignment of MoBre2 from *M. grisea*, yBre2 from *S. cerevisiae* and ASH2L from mammals. Numbers refer to amino acid residues. Identical residues among these proteins are shaded in black, whereas similar residues are shaded in grey. Panel D. Alignment of the conserved domains of MoSdc1 from *M.grisea*, ySdc1 from *S. cerevisiae* and DPY30 from mammals. Panel E. Alignment of the conserved domains of MoSpp1 from *M. oryzae*, Cps40 from *S. cerevisiae* and DPY30 from mammals.

**Fig. S2.**
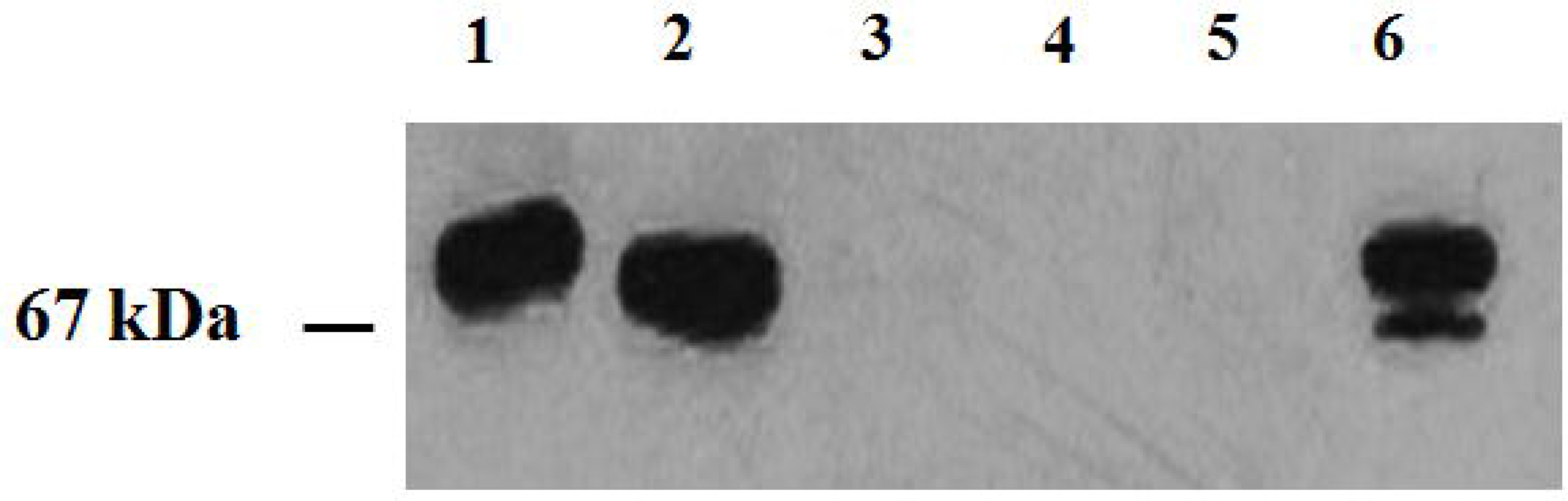
Identification of 3 × MoBre2-Flag mutant strains by western blotting. Immunoblotting with anti-FLAG antibody. Lanes 1 to 5 are five transformants, and lane 6 is a positive control.

**Fig. S3.**
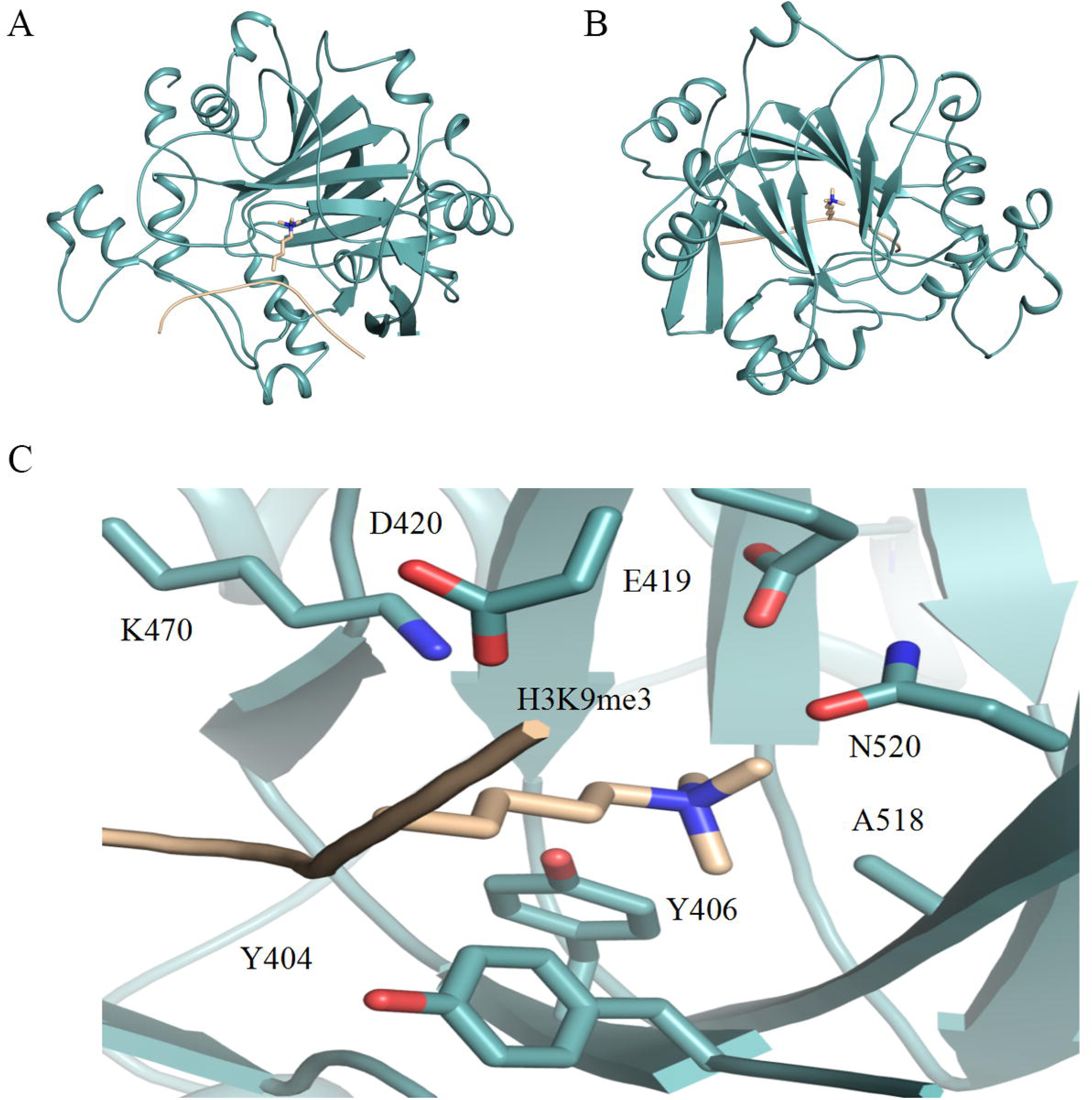
Modelled structure of the core tudor domain of MoJmjd2 in complex with histone H3K9me3. Shown are side view (A), the key residues caging the trimethylated H3K27 (C); and top view (B) a modelled MoJmjd2 (cyan)- H3K27me3 (gold) complex.

**Fig. S4.**
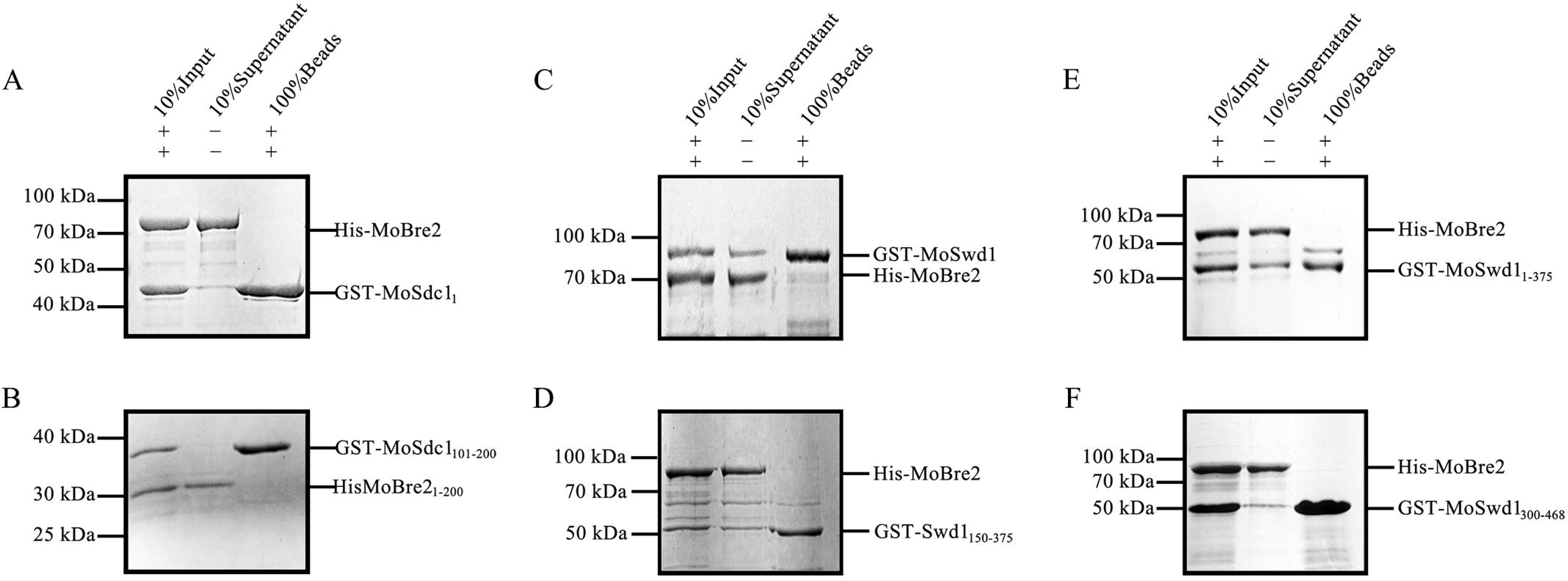
GST-tag-based pull-down assay of MoBre2 and MoSwd1. Panel A. Full-length MoBre2 cannot interact with the N-terminal region of MoSdc1 (residues 1-100). Panel B. The fragments of MoSdc1 (residues 101-200) cannot interact with MoBre2_1-200_. Panel C. GST-based pull-down assay with MoBre2 and MoSwd1. GST-tagged MoSwd1 was incubated with His-tagged MoBre2. The eluate was subjected to SDS-PAGE. Panels D-F. Full-length MoBre2 interacts with the three deletion mutants of MoSwd1_1-375_ (residues 1-375), MoSwd1_150-375_ (residues 150-375) and MoSwd1_375-468_ (residues 375-468).

**Fig. S5.**
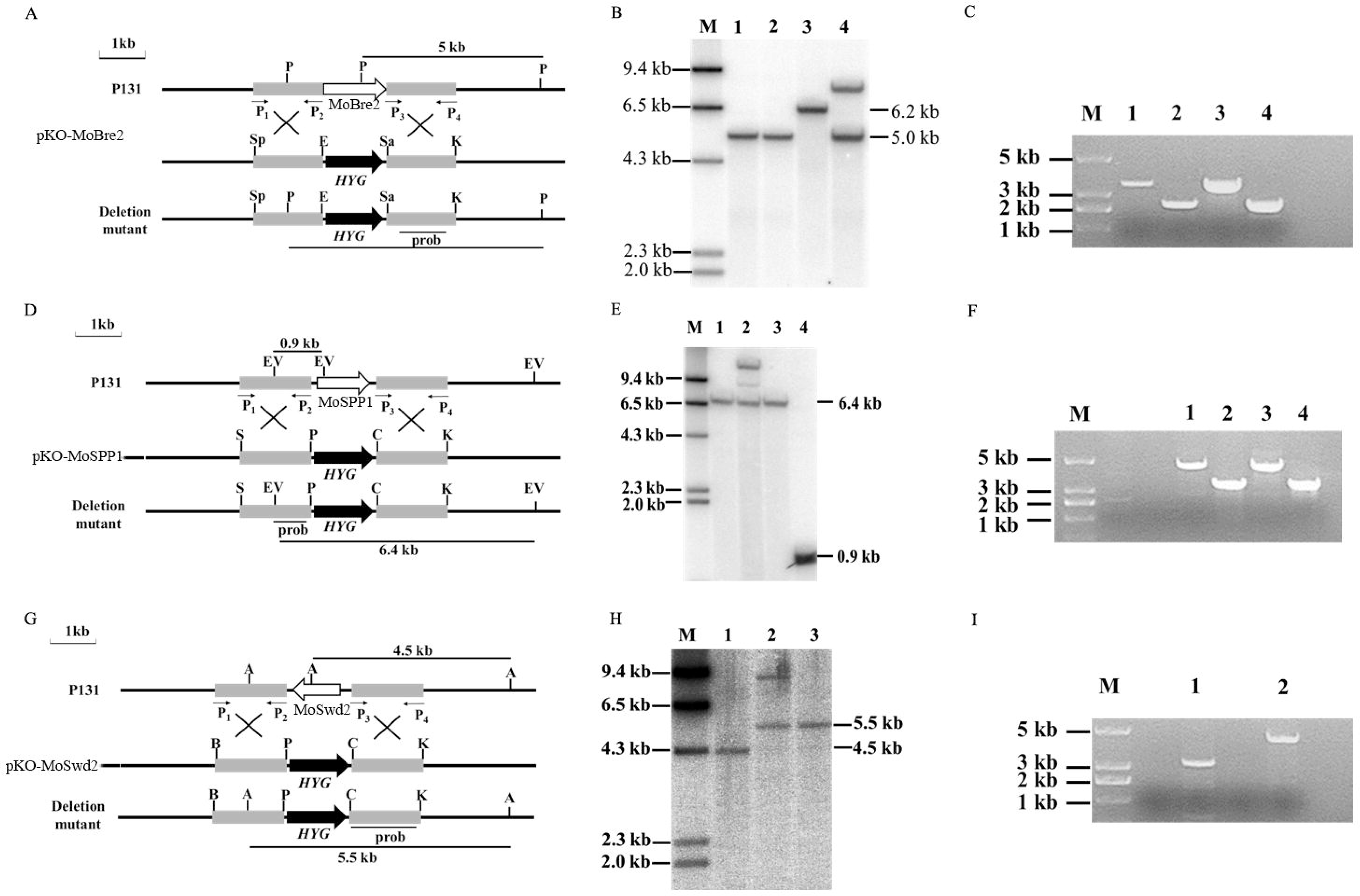
Schematic diagram of the deletion strategies for *MoBre2*, *MoSpp1* and *MoSwd2*. Panel A. Schematic diagram of the *MoBre2* deletion strategy. P, Sp, E, Sa, and K indicate the recognition sites of *Pst* I*, Spe* I*, Eco* RI*, Sal* I, and *Kpn* I, respectively. HYG, hygromycin phosphotransferase gene. P1 to P4 are the PCR primers *MoBre2*LF, *MoBre2*LR, *MoBre2*RF, and *MoBre2*RR. Panel B. Southern blot analysis of the *MoBre2* deletion mutants. *Pst* I-digested genomic DNA was hybridized with the probe (marked in A). Lane M, *HindIII*-digested fragments of λDNA as markers; lane 1, wild-type strain P131; lanes 2 to 4, *MoBre2* deletion mutants Δ*mobre2*KO1, Δ*mobre2*KO2, and Δ*mobre2*KO3. **C.** The Δ*mobre2*KO2 deletion mutant was verified with the primer pairs *MoBre2*F/HYBLBCK and *MoBre2*R/HYBRBCK. Lane M, marker; lane 1, Δ*mobre2*KO2 amplification product with *MoBre2*F/HYBLBCK; lane 2, Δ*mobre2*KO2 amplification product with *MoBre2*R/HYBRBCK; lanes 3 and 4 are repeats of lanes 1 and 2. Panel D. Schematic diagram of the *MoSpp1* deletion strategy. EV, S, P, C, and K indicate the recognition sites of *Eco*RV*, Spe*I*, Pst*I, *Cla* I,and *Kpn* I, respectively. HYG, hygromycin phosphotransferase gene. P1 to P4 are the PCR primers *MoSpp1*LF, *MoSpp1*LR, *MoSpp1*RF, and *MoSpp1*RR. Panel E. Southern blot analysis of the Δ*mospp1* deletion mutants. *Eco*RV-digested genomic DNA wash ybridized with the probe (marked in D). Lane M, HindIII-digested fragments of λDNA as markers; lanes 1 to 3, *MoSpp1* deletion mutants Δ*mospp1*KO3, Δ*mospp1*KO2, and Δ*mospp1*KO1; lane 4, wild-type strain P131. Panel F. The Δ*mospp1*KO1 and Δ*mospp1*KO3 deletion mutants were verified by PCR with the primer pairs *MoSpp1*F/HYBLBCK and *MoSpp1*R/HYBRBCK. Lane M, marker; lane 1, Δ*mospp1*KO1 amplification product with *MoSpp1*F/HYBLBCK; lane 2, Δ*mospp1*KO1 amplification product with *MoSpp1*R/HYBRBCK; lane 3, Δ*mospp1*KO3 amplification product with *MoSpp1*F/HYBLBCK; lane 4, Δ*mospp1*KO3 amplification product with *MoSpp1*R/HYBRBCK. Panel G. Schematic diagram of the Δ*moswd2* deletion strategy. A, B, P, C, and K indicate the recognition sites of *Apa* I*, Bam* HI*, Pst* I*, Cla* I, and *Kpn*, respectively. HYG, hygromycin phosphotransferase gene. P1 to P4 are the PCR primers *MoSwd2*LF, *MoSwd2*LR, *MoSwd2*RF, and *MoSwd2*RR. Panel H. Southern blot analysis of the *MoSwd2* deletion mutants. *Apa* I-digested genomic DNAwas hybridized with the probe (marked in G). Lane M, *Hind* III-digested fragments of λDNA as markers; lane 1, wild-type strain P131; lanes 2 and 3, Δ*moswd2* deletion mutants Δ*moswd2*KO1 and Δ*moswd2*KO2. Panel I.The Δ*moswd2*KO2 deletion mutant was verified by PCR with the primer pairs *MoSwd2*F/HYBLBCK and *MoSwd2*R/HYBRBCK. Lane M, marker; lane 1, Δ*moswd2*KO2 amplification product with *MoSwd2*F/HYBLBCK; lane 2, Δ*moswd2*KO2 amplification product with *MoSwd2*R/HYBRBCK.

**Fig. S6.**
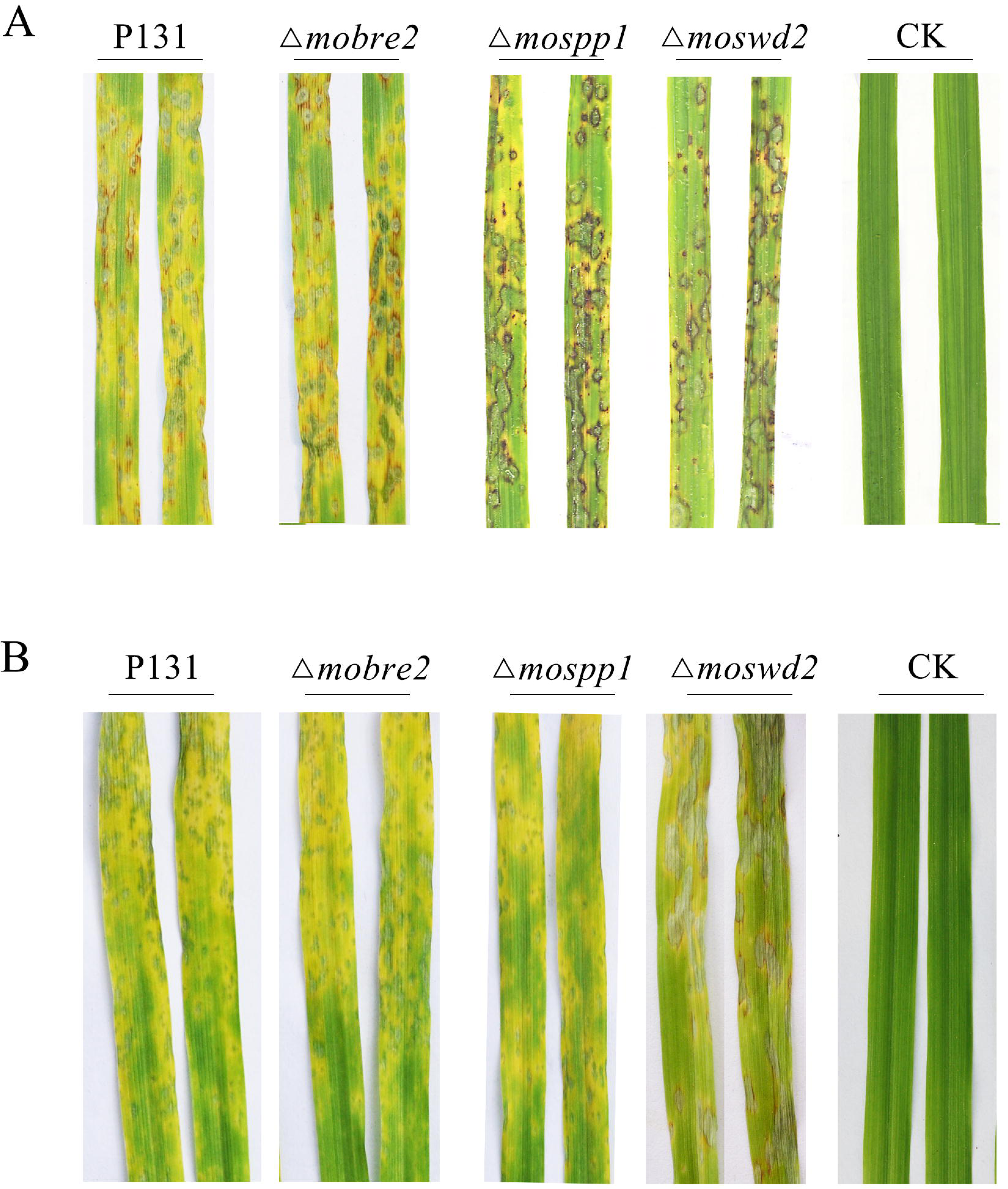
Inoculation test of the Δ*mobre2*, Δ*mospp1* and Δ*moswd2* deletion mutants. Rice leaves were sprayed with conidial suspensions (1 × 10^5^ spores/mL) of the wild-type P131, or of the deletion mutants Δ*mobre2*, Δ*mospp1* and Δ*moswd2*. The inoculated leaves were photographed 5 d post inoculation.

**Fig. S7.**
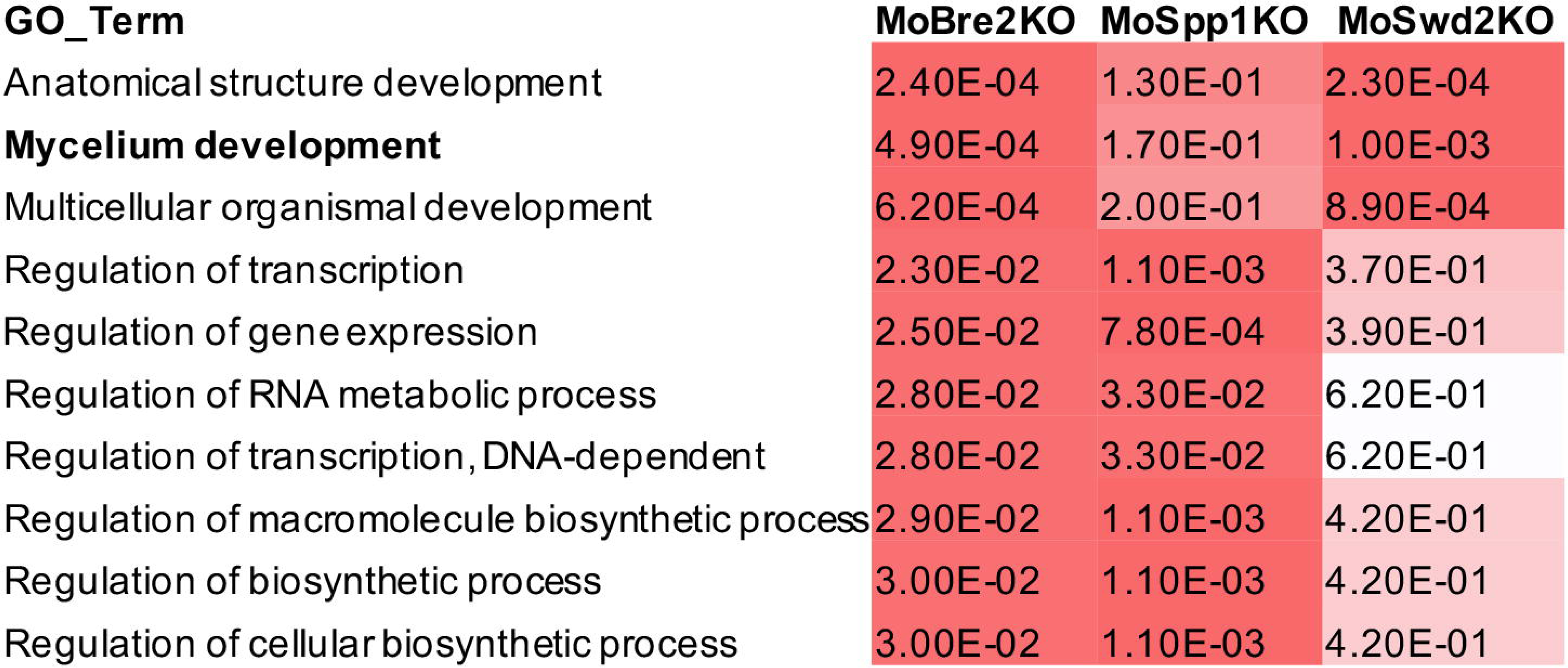
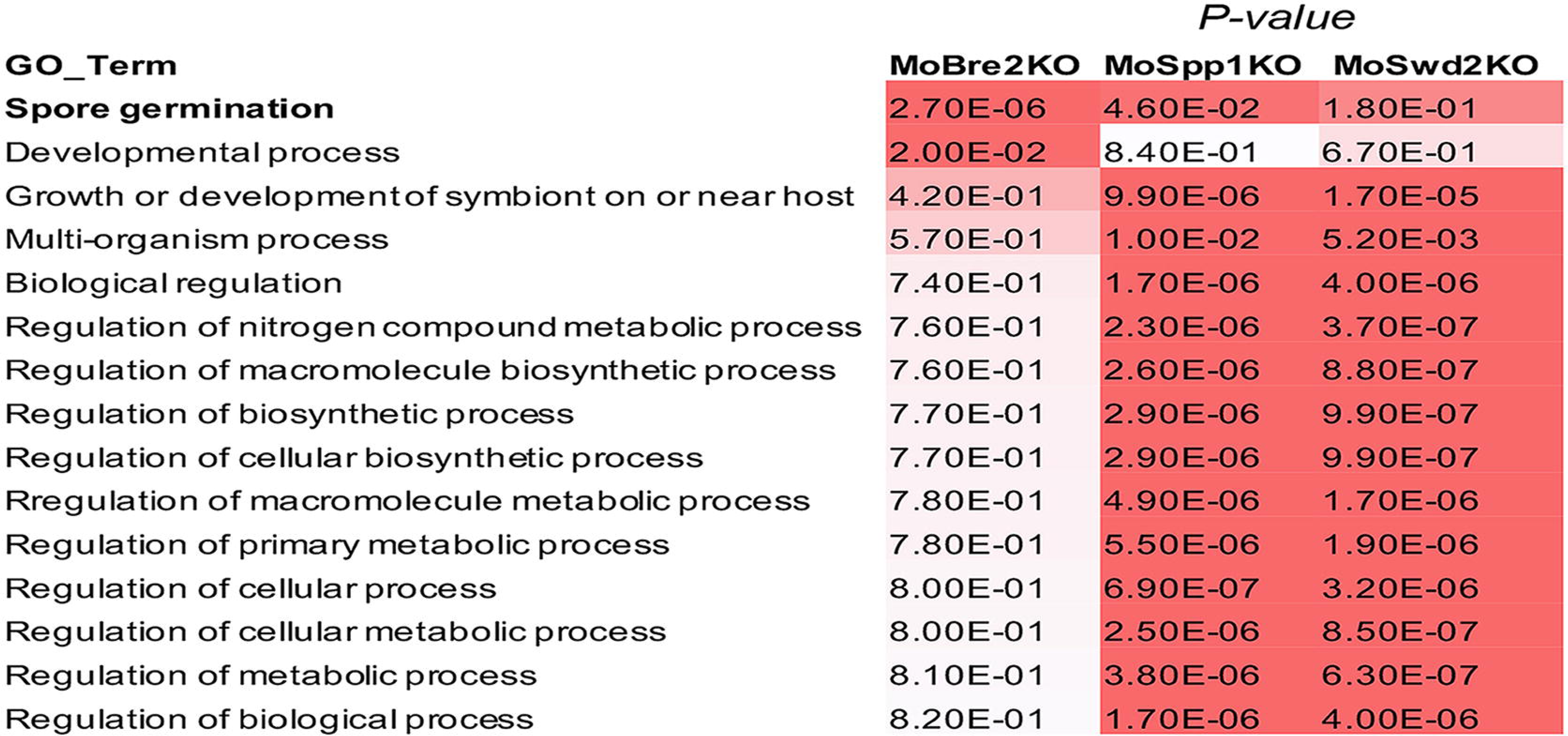
GO term analysis of the candidate genes involved in mycelium development with H3K4me3-ChIP-seq peaks decreased in Δ*mobre2*, Δ*mospp1* and Δ*moswd2* mutants. The value shown on the table represents the frequency of the GO term in the Gene Ontology annotation database. The colour represents the enrichment p-value (Fisher’s exact test) of each GO term. The scaled relative color are shown in the heatmap.

**Table S1.** Names and sequences of the 29 COMPASS-like proteins from *Magnaporthe oryzae*, *Saccharomyces cerevisiae*, *Fusarium graminearum*, *Arabidopsis*, *Drosophila*, and mammals shown in Figure 1.

**Table S2.** Tandem mass spectrometry analysis to identify five subunits of COMPASS-like proteins.

**Table S3.** Primers used in this study.

**Table S4.** List of the hypothetical genes with decreased H3K4me3-ChIP-seq peaks compared with the wild type strain in the Δ*mobre2*, Δ*mospp1* or Δ*moswd2* mutants.

**Table S5.** Gene Ontology analysis of the hypothetical genes with decreased H3K4me3-ChIP-seq peaks compared with the wild type strain in the Δm*obre2*, Δ*mospp1* or Δ*moswd2* mutants.

**Table S6.** Gene Ontology analysis of the pathogenesis-related genes with decreased H3K4me3-ChIP-seq peaks in the Δ*mobre2*, Δ*mospp1* or Δ*moswd2* mutants.

**Table S7.** List of the hypothetical genes decreased in H3K4me3-ChIP-seq levels overlapped with downregulated expression genes in RNA-seq in the Δ*mobre2*, Δ*mospp1* or Δ*moswd2* mutants.

**Table S8.** GO term analysis of candidate genes involved in spore germination with decreased in both H3K4me3-ChIP-seq signals and RNA-seq signals in the Δ*mobre2*, Δ*mospp1* or Δ*moswd2* mutants.

**Table S9.** List of genes decreased in H3K4me3-ChIP-seq levels and downregulated expression in RNA-seq in the Δ*mobre2*, Δ*mospp1* or Δ*moswd2* mutants.

## References

Agger, K., Cloos, P. A., Christensen, J., Pasini, D., Rose, S., Rappsilber. J., Issaeva, I., Canaani, E., Salcini, A. E. and Helin, K. (2007). UTX and JMJD3 are histone H3K27 demethylases involved in HOX gene regulation and development. Nature 449, 731–734.

Agger, K., Nishimura, K., Miyagi, S., Messling, J. E., Rasmussen, K. D. and Helin, K. (2019). The KDM4/JMJD2 histone demethylases are required for hematopoietic stem cell maintenance. Blood 134(14), 1154–1158.

Ali, A. and Tyagi, S. (2017). Diverse roles of capitalise acronyms as in the original complex in the functions of the Set1 histone methyltransferase family. Journal of Biosciences 42, 155–159.

Allis, C. D., Berger, S. L., Cote, J., Dent, S., Jenuwien, T., Kouzarides, T., Pillu, L., Reinberg, D., Shi, Y., Shiekhattar, R. et al. (2007). New nomenclature for chromatin-modifying enzymes. Cell 131, 131–633.

Amar, M. S., Sun, Y. H., Li, L., Zhang, W. J., Wu, T. M., Zhao, S. Y., Qin, Z. H. and Dalton, S. (2015). Cell-cycle control of bivalent epigenetic domains regulates the exit from pluripotency. Stem Cell Reports 5, 1–14.

Ardehali, M. B., Mei, A., Zobeck, K. L., Caron, M., Lis, J. T. and Kusch, T. (2011). *Drosophila* Set1 is the major histone H3 lysine 4 trimethyltransferase with role in transcription. The EMBO Journal 30, 2817–2828.

Bachleitner, S., Sørensen, J. L., Matthews, A. G., Sulyok, M., Studt, L. and Strauss, J. (2019). Evidence of a demethylase-independent role for the H3K4-specific histone demethylases in *Aspergillus nidulans* and *Fusarium graminearum* secondary metabolism. Frontiers in Microbiology 10, 1759.

Berger, S. L. (2007). The complex language of chromatin regulation during transcription. Nature 447, 407–412.

Bernstein, B. E., Mikkelsen, T. S., Xie, X., Kamal, M., Huebert, D. J., Cuff, J., Fry, B., Meissner, A., Wernig, M., Plath, K. et al. (2006). A bivalent chromatin structure marks key developmental genes in embryonic stem cells. Cell 125, 315–326.

Bhaumik, S. R., Smith, E. and Shilatifard, A. (2007). Covalent modifications of histones during development and disease pathogenesis. Nature Structural and Molecular Biology 14, 1008–1016.

Cao, Z. J., Yin, Y., Sun, X., Han, J., Sun, Q. P., Lu, M., Pan, J. B. and Wang, W. X. (2016). An ash1-like protein MoKMT2H null mutantis delayed for conidium germination and pathogenesis in *Magnaporthe oryzae*. BioMed Research International 10, 1575–1585.

Chen, Y., Cao, F., Wan, B., Dou, Y. and Lei, M. (2012). Structure of the SPRY domain of human ASH2L and its interactions with RbBP5 and DPY30. Cell Research 22, 598–602.

Cho, Y. W., Hong, T., Hong, S., Guo, H., Yu, H., Kim, D., Guszczynski, T., Dressler, G. R., Copeland, T. D., Kalkum, M. et al. (2007). PTIP associates with MLL3-and-MLL4-containing histone H3 lysine 4 methyltransferase complex. The Journal of Biological Chemistry 282, 20395–406.

Connolly, L. R, Smith, K. M. and Freitag, M. (2013). The *Fusarium graminearum* histone H3K27 methyltransferase KMT6 regulates development and expression of secondary metabolite gene clusters. PloS Genetics 9, e1003916.

Czermin, B., Melfi, R., McCabe, D., Seitz, V., Imhof, A. and Pirrotta, V. (2002). *Drosophila* enhancer of zeste/esc complexes have a histone H3 methyltransferase activity that marks chromosomal polycomb sites. Cell 111, 185–196.

Dang, J. L. and Jones, J. D. G. (2001). Plant pathogens and integrated defence response to infection. Nature 411, 826–833.

Deng, S., Gu, Z., Yang, N., Li, L., Yue, X., Que, Y., Sun, G., Wang, Z. and Wang, J. (2016). Identification and characterization of the peroxin 1 gene *MoPEX1* required for infection-related morphogenesis and pathogenicity in *Magnaporthe oryzae*. Scientific Reports 10, 1038–36292.

Dover, J., Schneider, J., Tawiah-Boateng, M. A., Wood, A., Dean, K., Johnston, M. and Shilatifard, A. (2002). Methylation of histone H3 by COMPASS requires ubiquitination of histone H2B by Rad6. Journal of Biological chemistry 277, 28368–28371.

D’Urso, A., Takahashi, Y., Xiong, B., Marone, J., Coukos, R., Hinchliff, C. R., Wang, J. P., Shilatifard, A. and Brickner, J. H. (2016). Set1/COMPASS and Mediator are repurposed to promote epigenetic transcriptional memory. eLife 5, e1669.

Ernst, P. and Vakoc, C. R. (2012). WRAD: enabler of the Set1-family of H3K4 methyltransferases. Brief Funct Genomics 11, 217–226.

Hsu, P. L., Li, H., Lau, H. T., Ong, S. E., Chatterjee, C. and Zheng, N. (2018). Crystal structure of the COMPASS H3K4 methyltransferase catalytic module. Cell 174, 1106–1116.

Huh, A., Dubey, A., Kim, K., Jeon, J. and Lee, Y. H. (2017). MoJMJ1, Encoding a histone demethylase containing JmjC domain, is required for pathogenic development of the rice blast fungus, *Magnaporthe oryzae*. Plant Pathol. J. 33(2), 193–205.

Jiang, D. H., Kong, N. C., Gu, X. F., Li, Z. C. and He, Y. H. (2011). Arabidopsis COMPASS-like complexes mediate histone H3 lysine-4 trimethylation to control floral transition and plant development. PLoS Genetics 7, e1001330.

Kanda, H., Nguyen, A., Chen, L., Okano, H. and Hariharana, I. K. (2013). The *Drosophila* ortholog of *MLL3* and *MLL4, trithorax related*, functions as a negative regulator of tissue growth. Molecular and Cellular Biology 33, 1702–1710.

Kelley, L. A., Mezulis, S., Yates, C. M., Wass, M. N. and Sternberg, M. J. (2015). The phyre2web portal for protein modeling, prediction and analysis. Nature Protocols 10, 845–858.

Kim, H. J., Han, J. H., Kim, K. S. and Lee, Y. H. (2014). Comparative functional analysis of the velvet gene family reveals unique roles in fungal development and pathogenicity in *Magnaporthe oryzae*. Fungal Genetics and Biology 66, 33–43.

Kornberg, R. D. (1974). Chromatin structure: a repeating unit of histones and DNA. Science 184, 868–871.

Kouzarides, T. (2007). Chromatin modifications and their function. Cell 128, 693–705.

Krogan, N. J., Dover, J., Wood, A., Schneider, J., Heidt, J., Boateng, M. A., Dean, K., Ryan, O. W., Golshani, A., Johnston, M. et al. (2003). The Paf1 complex is required for histone H3 methylation by COMPASS and Dot1p: linking transcriptional elongation to histone methylation. Molecular Cell 11, 721–729.

Krogan, N. J., Kim, M., Tong, A., Golshani, A., Cagney, G., Canadien, V., Richards, D. P., Beattie, B. K., Emili, A., Boone, C. et al. (2003). Methylation of histone H3 by Set2 in *Saccharomyces cerevisiae* is linked to transcriptional elongation by RNA polymerase II. Molecular & Cellular Biology 23, 4207–4218.

Lachner, M. and Jenuwein, T. (2002). The many faces of histone lysine methylation. Current Opinion in Cell Biology 14, 286–298.

Lee, J. H. and Skalnik, D. G. (2008). Wdr82 is a C-terminal domain-binding protein that recruits the Setd1A Histone H3-Lys4 methyltransferase complex to transcription start sites of transcribed human genes. Molecular and Cellular Biology 28, 609–18.

Li, Y., Han, J., Zhang, Y., Cao, F., Liu, Z., Li, S., Wu, J., Hu, C., Wang, Y., Shuai, J. et al. (2016). Structural basis for activity regulation of MLL family methyltransferases. Nature 530, 447–452.

Liu, C. Y., Li, Z. G., Xing, J. J., Yang, J., Wang, Z., Zhang, H., Chen, D., Peng, Y. L. and Chen, X. L. (2018). Global analysis of sumoylation function reveals novel insights into development and appressorium-mediated infection of the rice blast fungus. New Phytologist 219, 1031–1047.

Liu, W., Xie, S., Zhao, X., Chen, X., Zheng, W., Lu, G., Xu, J. R. and Wang, Z. (2010). A homeobox gene is essential for conidiogenesis of the rice blast fungus *Magnaporthe oryzae*. Molecular Plant-Microbe Interactions 23, 366–375.

Liu, Y., Liu, N., Yin, Y. N., Chen, Y., Jiang, J. H. and Ma, Z. H. (2015). Histone H3K4 methylation regulates hyphal growth, secondary metabolism and multiple stress responses in *Fusarium graminearum*. Environmental Microbiology 17, 4615–4630.

Luger, K., Maeder, A. W., Richmond, R. K., Sargent, D. F. and Richmond, T. J. (1997). Crystal structure of the nucleosome core particle at 2.8-angstrom resolution. Nature 389, 251–260.

Mao, Z., Pan, L., Wang, W. X., Sun, J., Shan, S., Dong, Q., Liang, X. P., Dai, L. C., Ding, X. J., Chen, S. et al. (2014). Anp32e, a higher eukaryotic histone chaperone directs preferential recognition for H2A.Z. Cell Research 24, 389–399.

Matthews, A. G., Berger, H., Sasaki, T., Wittstein, K., Gruber, C., Lewis, Z. A. and Strauss, J. (2016). KdmB, a Jumonji histone H3 demethylase, regulates genome-wide H3K4 trimethylation and is required for normal induction of secondary metabolism in *Aspergillus nidulans*. PLoS Genetics 12(8), e1006222.

Matthews, A. G., Noble, L. M., Gruber, C., Berger, H., Sulyok, M., Marcos, A. T., Joseph Strauss, J. and Andrianopoulos, A. (2015). KdmA, a histone H3 demethylase with bipartite function, differentially regulates primary and secondary metabolism in *Aspergillus nidulans*. Molecular Microbiologyl 96(4), 839–860.

Miller, T., Krogan, N. J., Dover, J., Erdjument, B. H., Tempst, P., Johnston, M., Greenblatt, J. F. and Shilatifard, A. (2001). COMPASS: a complex of proteins associatedwith a trithorax-related SET domain protein. Proceedings of the National Academy of Sciences of the United States of America 98, 12902–12907.

Mohan, M., Herz, H-M., Smith, E. R., Zhang, Y., Jackson, J., Washburn, M. P., Florens, L., Eissenberg, J. C. and Shilatifard, A. (2011). The COMPASS family of H3K4 methylases in *Drosophila*. Molecular and Cell-ular Biology 31, 4310–4318.

Ng, H. H., Robert, F., Young, R. A. and Struhl, K. (2003). Targeted recruitment of set1 histone methylase by elongating Pol II provides a localized mark and memory of recent transcriptional activity. Molecular Cell 11, 709–719.

Ng, S. S., Kavanagh, K. L., Mcdonough, M. A., Butler, D., Pilka, E. S., Lienard, B. M., Bray, J. E., Savitsky, P., Gileadi, O., von Delft, F. et al. (2007). Crystal structures of histone demethylase JMJD2A reveal basis for substrate specificity. Nature 448, 87–91.

Palmer, J. M., Bok, J. W., Lee, S., Dagenais, T. R. T., Andes, D. R., Kontoyiannis, D. P. and Keller, N. P. (2013). Loss of *CclA*, required for histone 3 lysine 4 methylation, decreases growth but increases secondary metabolite production in *Aspergillus fumigatus*. PeerJ 1, e4.

Patkar, R. N., Ramospamplona, M., Gupta, A. P., Fan, Y. and Naqvi, N. I. (2012). β-oxidation regulates organellar integrity and is necessary for conidial germination and invasive growth in *Magnaporthe oryzae*. Molecular Microbiology 86, 1345–1363.

Pham, K. T. M., Inoue, Y., Vu, B. V., Nguyen, H. H., Nakayashiki, T., Ikeda, K. and Nakayashiki, H. (2015). MoSet1 (histone H3K4 methyltransferase in *Magnaporthe oryzae*) regulates global gene expression during infection-related morphogenesis. PLoS Genetics 11, e1005385.

Rahman, H. and De Sousa, R. D. (2014). Role of MoAnd1-mediated nuclear positioning in morphogenesis and pathogenicity in the rice blast fungus, Magnaporthe oryzae. Fungal Genetics & Biology 69, 43–51.

Robert, X. and Gouet, P. (2014). Deciphering key features in protein structures with the new endscript server. Nucleic Acids Research 42, W320–W324.

Roguev, A., Schaft, D., Shevchenko, A., Pijnappel, W. W., Wilm, M., Aasland, R. and Stewart, A. F. (2001). The *Saccharomyces cerevisiae* Set1 complex includes an Ash2 homologue and methylates histone 3 lysine 4. EMBO Journal 20, 7137–7148.

Shilatifard, A. (2008). Molecular implementation and physiological roles for histone H3 lysine 4 (H3K4) methylation. Current Opinion in Cell Biology 20, 341–348.

Shilatifard, A. (2012). The COMPASS family of histone H3K4 methylases: mechanisms of regulation in development and disease pathogenesis. Annual Review of Biochemistry 81, 65–95.

Shinohara, Y., Kawatani, M., Futamura, Y., Osada, H. and Koyama, Y. (2016). An overproduction of astellolides induced by genetic disruption of chromatinremodeling factors in Aspergillus oryzae. J. Antibiot. 69, 4–8.

Smith, E. and Shilatifard, A. (2010). The chromatin signaling pathway: diverse mechanisms of recruitment of histone-modifying enzymes and varied biological outcomes. Molecular Cell 40, 689–701.

Smith, E. R., Lee, M. G., Winter, B., Droz, N. M., Eissenberg, J. C., Shiekhattar, R. and Shilatifard, A. (2008). Drosophila UTX is a histone H3 Lys27 demethylase that colocalizes with the elongating form of RNA polymerase II. Molecular & Cellular Biology 28, 1041–1046.

Stassen, M. J., Bailey, D., Nelson, S., Chinwalla, V. and Harte, P. J. (1995). The *Drosophila* trithorax proteins contain a novel variant of the nuclear receptor type DNA binding domain and an ancient conserved motif found in other chromosomal proteins. Mechanisms of Development 52, 209–23.

Strath, B. D. and Allis, C. D. (2000). The language of covalent histone modifications. Nature 403, 41–45.

Studt, L., Rösler, S. M., Burkhardt, I., Arndt, B., Freitag, M., Humpf, H. U., Dickschat, J. S. and Tudzynsji, B. (2016). Knock-down of the methyltransferase Kmt6 relieves H3K27me3 and results in induction of cryptic and otherwise silent secondary metabolite gene clusters in *Fusarium fujikuroi*. Environ. Microbiol. 18, 4037–4054.

Takahashi, Y. H., Westfield, G. H., Oleskie, A. N., Trievel, R. C., Shilatifard, A. and Skiniotis, G. (2011). Structural analysis of the core COMPASS family of histone H3K4 methylases from yeast to human. Proceedings of the National Academy of Sciences of the United States of America 108, 20526–20531.

Tremblay V., Zhang P., Chaturvedi C. P., Thornton J., Brunzelle J. S., Skiniotis G., Shilatifard A., Brand M. and Counture J. F. (2014). Molecular basis for DPY-30 association to COMPASS-like and NURF complexes. Structure 22, 1821–1830.

Tschiersch, B., Hofmann, A., Krauss, V., Dorn, R., Korge, G. and Reuter, G. (1994). The protein encoded by the *Drosophila* position-effect variegation suppressor gene Su(var)3–9 combines domains of antagonistic regulators of homeotic gene complexes. EMBO Journal 13, 3822–3831.

Wang, G. H., Wang, C. F., Hou, R., Zhou, X. Y., Li, G. T., Zhang, S. J. and Xu, J. R. (2012). The AMT1 arginine methyltransferase geneis important for plant infection and normal hyphal growth in *Fusarium graminearum*. PLoS One 7(5), e38324.

Wang, J. K., Tsai, M. C., Poulin, G., Adler, A. S., Chen, S., Liu, H., Shi, Y. and Chang, H. Y. (2010). The histone demethylase UTX enables RB-dependent cell fate control. Genes & Development 24, 327–332.

Wang, W. X., Chen, Z., Mao, Z., Zhang, H. H., Ding, X. J., Chen, S., Zhang, X. D., Xu, R. M. and Zhu, B. (2011). Nucleolar protein Spindlin1 recognizes H3K4 methylation and stimulates the expression of rRNA genes. EMBO Reports 12, 1160–1166.

Workman, J. L. and Kingston, R. E. (2003). Alteration of nucleosome structure as a mechanism of transcriptional regulation. Annual Review of Biochemistry 67, 545–579.

Yan, X. and Talbot, N. J. (2016). Investigating the cell biology of plant infection by the rice blast fungus *Magnaporthe oryzae*. Current Opinion in Microbiology 34, 147–153.

Yang, N., Wang, W. X., Wang, Y., Wang, M. Z., Zhao, Q., Rao, Z. H., Zhu, B. and Xu, R. M. (2012). Distinct mode of methylated lysine-4 of histone H3 recognition by tandem tudor-like domains of Spindlin1. Proceedings of the National Academy of Sciences of the United States of America 109, 17954–17959.

Kim, Y. N., Park, S-Y, Kim, D. Y., Choi, J. Y., Lee, Y-H, Lee, J-H and Choi, W. B. (2013). Genome-scale analysis of ABC transporter genes and characterization of the ABCC type transporter genes in *Magnaporthe oryzae*. Genomics 10, 354–361.

Kou, Y. J., Tan, Y. H., Ramanujam, R. I. and Naqvi, N. (2016). Structure-function analyses of the Pth11 receptor reveal an important role for CFEM motif and redox regulation in rice blast. New Phytologist 214, 330–342.

Zhang, H., Zhao, Q., Guo, X., Guo, M., Qi, Z., Tang, W., Dong, Y., Ye, W., Zheng, X. and Wang, P. (2014). Pleiotropic function of the putative zinc-finger protein MoMsn2 in *Magnaporthe oryzae*. Molecular Plant-Microbe Interactions 27, 446–460.

Zhang, M. Y., Sun, X., Cui, L. E., Yin, Y., Zhao, X. Y., Pan, S. and Wang, W. X. (2018). The plant infection test: spray and wound-mediated inoculation with the plant pathogen *Magnaporthe grisea*. Journal of Visualized Experiments 138, 10.3791/57675.

